# Neuronal APOE4 drives damaging lipid accumulation via contact-dependent neuron-oligodendrocyte-microglia interaction in Alzheimer’s disease

**DOI:** 10.64898/2025.12.04.692390

**Authors:** Min Joo Kim, Jessica Blumenfeld, Yaqiao Li, Oscar Yip, Luokang Yao, Stephanie Ancheta, Samuel De Leon, Zoe Platow, Zherui Liang, David Shostak, Kaylie Suan, Yanxia Hao, Nicole Koutsodendris, Claire Ellis, Jeremy Nguyen, Yadong Huang

## Abstract

Apolipoprotein E4 (APOE4) confers the greatest genetic risk for developing Alzheimer’s disease (AD). With APOE4 broadly expressed in the brain, its cell-type-specific roles in AD pathogenesis are only beginning to be defined. Here, we show that neuronal APOE4 expression drives damaging lipid accumulation in hippocampal neurons, oligodendrocytes, and microglia, with preferential buildup of peroxidized lipids in microglia in a tauopathy mouse model. Neuron-specific removal of APOE4 abolished this lipid phenotype, whereas neuron-specific expression of APOE4 was sufficient to recapitulate it, demonstrating that neuronal APOE4 is both necessary and sufficient for lipid accumulation. Strikingly, the association between lipid burden, microgliosis, and neurodegeneration was strongest in mice with neuron-specific APOE4 expression. Single-nucleus RNA sequencing revealed neuronal APOE4-vulnerable neuron populations, as well as enrichment of disease-associated microglia and oligodendrocytes, all promoting lipid pathology. Primary mouse co-culture experiments showed that neuronal APOE4 drives microglial lipid accumulation via contact-dependent mechanisms involving uptake of lipids from neurons and oligodendrocytes. These findings establish neuronal APOE4 as a key driver of lipid accumulation via neuron-oligodendrocyte-microglia interactions, providing mechanistic insight into APOE4-driven lipid pathology in AD.

## Introduction

Alzheimer’s disease (AD) affects over 50 million people worldwide, with the apolipoprotein E4 (*APOE4*) gene as the strongest genetic risk factor for late-onset AD^1–3^. Carriers with two copies of the *APOE4* allele have a 10–15 fold increased risk of developing AD^4–6^, yet the multi-system cellular mechanisms underlying this vulnerability remain elusive. Functionally, APOE is primarily a lipid transporter^7,8^, with isoforms that differ in lipid-carrying and receptor-binding properties and thus contribute differentially to AD pathogenesis.

While APOE’s lipid carrier function has been well established, its role in lipid dysregulation and accumulation represents an understudied but potentially crucial aspect of AD pathogenesis^7,8^. Alois Alzheimer’s early descriptions of the disease noted lipid-laden glial cells^11^, and recent work has renewed interest in lipid metabolism dysfunction^9,10,12,13^. Lipid droplets (LDs) accumulate in aging and neurodegenerative brains^9,12,13^ serving as markers of cellular stress and potential contributors to disease progression. Increasing evidence implicates APOE4 as an important upstream regulator and casual driver of lipid and metabolic dysfunction in the brain, particularly within microglia^10,12,14,15^, highlighting APOE4 as a key target for understanding lipid dysregulation in AD pathogenesis. Beyond cell-autonomous metabolic defects, APOE4 also appears to disrupt neuron-glia lipid coupling^16^, impairing microglial clearance of excess neuronal lipids^12^. This maladaptive lipid exchange promotes LD buildup across cell types and fuels a cycle of glial dysfunction, positioning APOE4-driven lipid dysregulation as an important mechanism in AD pathogenesis^9,12,12,17,18^.

Despite substantial attention on microglia, the contribution of oligodendrocytes—one of the most lipid-rich and metabolically vulnerable cell types in the brain^19–21^—remains comparatively underexplored. Oligodendrocytes require tightly regulated lipid synthesis for myelin maintenance and are highly sensitive to metabolic and oxidative stress^19,21,22^, making them potential early responders to APOE4-driven lipid imbalance. Recent transcriptomic and mouse studies implicate oligodendrocyte dysfunction^23,24^, myelin instability^25^, and altered lipid processing^25,26^ in both aging and APOE4 contexts. Yet how oligodendrocytes interact with neurons and microglia to participate in, or contribute to, APOE4-dependent lipid dysregulation in AD remains largely unknown.

Growing evidence suggests that the cellular source of APOE4 expression critically influences AD progression^3^. While APOE4 is produced by multiple brain cell types—including astrocytes, microglia, neurons, and oligodendrocytes—neuron-derived APOE4 has garnered particular attention for its role in tau-mediated pathologies^23,27^. Neuron-specific APOE4 deletion reduces tau pathology and neurodegeneration^23^, while neuron-specific APOE4 expression is sufficient to recapitulate global APOE4 pathologies^27^, implicating neuronal APOE4 as a key pathogenic driver. However, the relationships between cell-type specific APOE4 expression, cellular lipid homeostasis, and neurodegeneration remains poorly characterized.

This study aims to define the role of neuronal APOE4 in lipid dysregulation across neurons and glia using a tauopathy mouse model expressing mutant human tau-P301S (PS19 line)^28^ and human APOE isoforms^23,27^. We show that neuronal APOE4 is both necessary and sufficient for driving LD and neutral lipid accumulation in oligodendrocytes and microglia via contact-dependent mechanisms involving microglial uptake of lipids from neurons and oligodendrocytes. These findings establish neuronal APOE4 as a key driver of microglial lipid accumulation through neuron-oligodendrocyte-microglia interactions, providing mechanistic insight into APOE4-driven lipid pathology in AD pathogenesis.

## Results

### PS19/E4 mice accumulate LDs in neurons and microglia in the hippocampus

To characterize how APOE isoforms differentially affect lipid profiles in the brain, we used previously generated mouse lines carrying a LoxP-floxed human *APOE4* or *APOE3* gene^29^. These floxed APOE-knock-in (E-KI) mice express homozygous human APOE4 or APOE3 in lieu of endogenous mouse ApoE. We crossbred these mice with a tauopathy line expressing the disease-causing human tau-P301S mutation on the 1N4R tau background (PS19 line)^28^. The resulting genotypes are referred to as PS19/E4 and PS19/E3 mice^23,24^.

To determine the effects of APOE isoforms on LDs, we evaluated mice at 10 months of age, when PS19 mice exhibit extensive hippocampal pathologies^28^. We first performed immunohistochemistry (IHC) with the LD-specific marker, perilipin-2 (PLIN2). Compared to PS19/E3 mice, PS19/E4 mice exhibited significantly more PLIN2^+^ LDs in both NeuN^+^ neurons (PLIN2^+^/NeuN^+^, neuronal LDs) and Iba1^+^ microglia (PLIN2^+^/Iba1^+^, microglial LDs) in the dentate gyrus (DG) of the hippocampus (Fig. 1a-c). In PS19/E4 or PS19/E3 hippocampi, LD levels in neurons or microglia did not significantly correlate with gliosis, AT8^+^ phosphorylated tau (p-tau), NeuN^+^ neuronal layer thickness, or hippocampal volume (Extended Data Fig. 1a-p), with two exceptions in PS19/E3 mice: microglial LDs correlated with hippocampal volume (Extended Data Fig. 1l), and neuronal LDs correlated with microgliosis (Extended Data Fig. 1m). These results suggest that LD accumulation in neurons or microglia may not contribute significantly or serve as strong predictors of tauopathy-related pathologies in PS19/E4 mice.

**Fig. 1.**
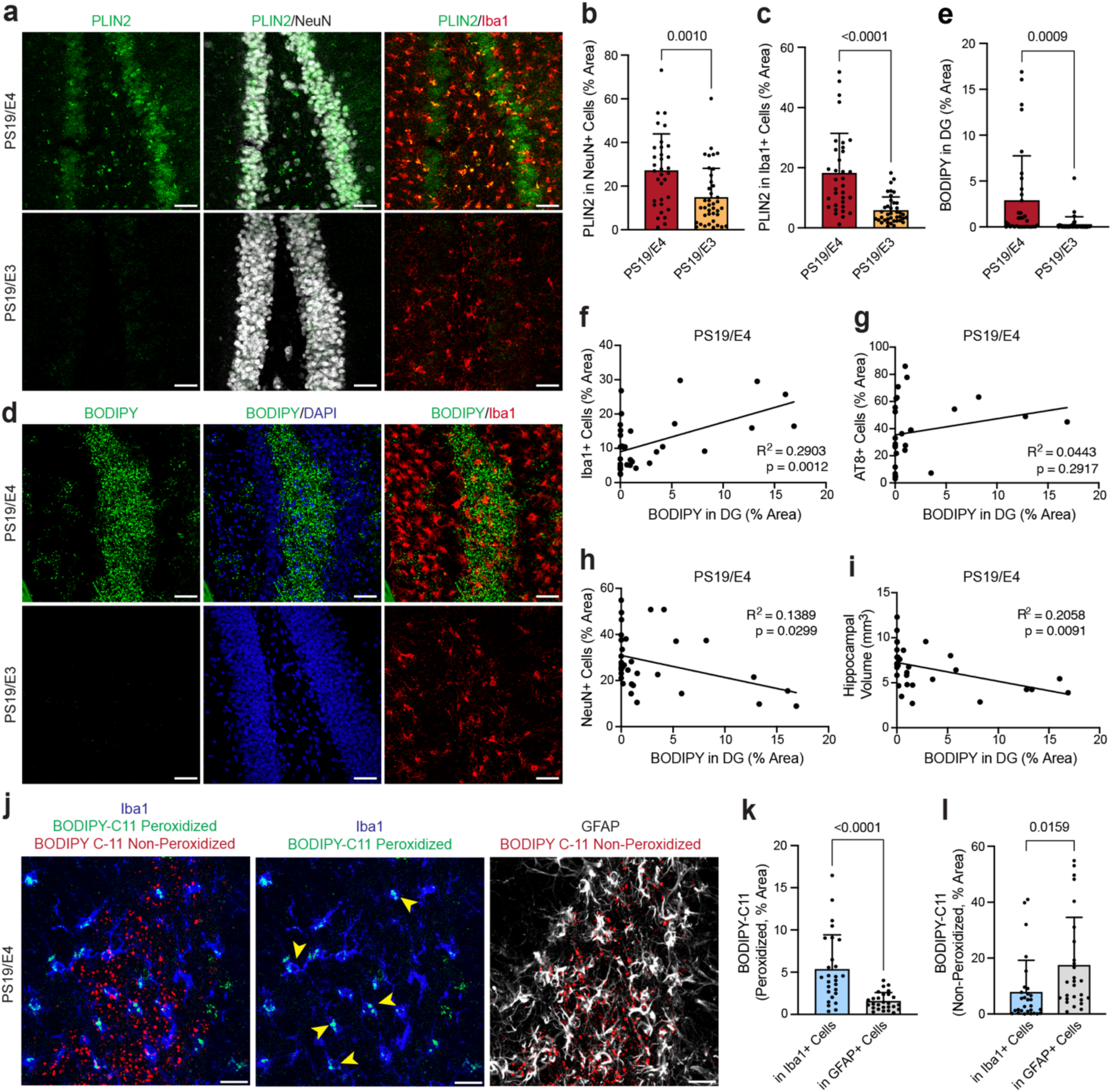
| APOE4 promotes lipid accumulation in the hippocampal dentate gyrus (DG) in PS19 tauopathy mice. **a,** Representative immunofluorescent images of PLIN2^+^ lipid droplets (green), NeuN^+^ neurons (white), and Iba1^+^ microglia (red) in the hippocampal DG of 10-month-old PS19/E4 and PS19/E3 mice. **b,** Quantification of percent PLIN2^+^ area coverage in NeuN^+^ neurons. **c,** Quantification of percent PLIN2^+^ area coverage in Iba1^+^ microglia. **d,** Representative immunofluorescent images of BODIPY^+^ neutral lipids (green), DAPI^+^ nuclei (white), and Iba1^+^ microglia (red) in the hippocampal DG of 10-month-old PS19/E4 and PS19/E3 mice. **a** and **d,** Scale bars, 40μm. **e,** Quantification of percent BODIPY^+^ area coverage in the hippocampal DG. **f-i,** Correlations between BODIPY^+^ % area in DG and Iba1^+^ % area (**f**), AT8^+^ % area (**g**), NeuN^+^ % area (**h**), and hippocampal volume (**i**) in PS19/E4 mice. **j,** Representative immunofluorescent images of BODIPY-C11^+^ peroxidized lipids (green), BODIPY-C11^+^ non-peroxidized lipids (red), Iba1^+^ microglia (blue), and GFAP^+^ astrocytes (white) in PS19/E4 mice. Scale bars, 20μm **k,** Quantifications of BODIPY-C11^+^ peroxidized lipids in Iba1^+^ cells and GFAP^+^ cells in PS19/E4 mice. **l,** Quantifications of BODIPY-C11^+^ non-peroxidized lipids in Iba1^+^ cells and GFAP^+^ cells in PS19/E4 mice. Quantified data in **b,c,e,k,l** are represented as mean ± s.e.m., unpaired two-sided t-test. For all experiments, PS19/E4, *n*= 35; and PS19/E3 *n*= 42. Pearson’s correlation analysis (two-sided) used for **f-i**.

Aging brains have been previously shown to accumulate LDs in microglia^9,17^. To assess aging effects on LD phenotypes in a tauopathy model expressing human APOE4, we quantified PLIN2^+^ LDs in PS19/E4 mice at different ages (∼ 4, 6, 8, and 10 months) (Extended Data Fig. 2a). We observed a linear increase in PLIN2^+^/NeuN^+^ neuronal LDs as mice age (Extended Data Fig. 2b). Interestingly, PLIN2^+^/Iba1^+^ microglial LDs exhibited an exponential increase beginning around 8 months of age, coinciding with the time when gliosis accelerates in PS19/E4 mice^28,21^ (Extended Data Fig. 2c). This trajectory suggests that in PS19/E4 mice, neurons accumulate LDs first in a steady, age-dependent manner, whereas microglial LD accumulation emerges later and increases sharply with onset of gliosis.

Together, these findings demonstrate that APOE4 expression promotes LD accumulation in both neurons and microglia in the hippocampus of PS19 tauopathy mice. Moreover, neuronal LD buildup appears earlier and progresses gradually with age, while microglial LD accumulation rises sharply during gliosis. These results support a model in which LDAMs emerge in response to age-dependent neuronal pathology in PS19/E4 hippocampi and contribute to the neuroinflammatory environment that further exacerbates neuronal stress.

### PS19/E4 mice accumulate neutral lipids in microglia and astrocytes in the hippocampus

Building on our analysis of LDs, we next examined whether neutral lipid accumulation showed a similar or distinct pattern. Interestingly, when we stained the hippocampus with BODIPY, a chemical dye specific for neutral lipids such as cholesterol esters and triglycerides, we observed a distribution that differed markedly from LDs. BODIPY^+^ neutral lipids were predominately localized to the DG and found mainly in Iba1^+^ microglia (Fig. 1d) and GFAP^+^ astrocytes (Extended Data Fig. 3a). Similar to LDs, however, the total percent area covered by the neutral lipid accumulation in the DG hilus of PS19/E4 mice was significantly greater than in PS19/E3 mice (Fig. 1e).

Neutral lipid accumulation was positively correlated with percent area coverage of Iba1^+^ microglia in PS19/E4 mice (Fig. 1f), indicating that neutral lipid burden increases with microgliosis. However, increased neutral lipids had no significant correlation with AT8^+^ p-tau percent area coverage (Fig. 1g). Conversely, the total neutral lipid area was negatively correlated with NeuN^+^ neuronal area and hippocampal volume, linking neutral lipid accumulation with neurodegeneration in PS19/E4 mice (Fig. 1h, i). Although overall neutral lipid coverage was low, PS19/E3 mice showed similar correlations with microgliosis and AT8^+^ area (Extended Data Fig. 3b,c) but did not exhibit the negative correlation with NeuN^+^ area and hippocampal volume observed in PS19/E4 mice (Extended Data Fig. 3d,e). We also found that neutral lipid accumulation followed a similar age-dependent trajectory as microglial LD accumulation, showing an exponential increase around the onset of gliosis at ∼8 months of age (Extended Data Fig. 3f, g).

### PS19/E4 mice preferentially accumulate peroxidized neutral lipids in microglia

To further characterize the type of neutral lipids accumulating and associated cellular regionality in PS19/E4 mouse brains, we stained with BODIPY 581/591C11 (BODIPY-C11), a reporter of lipid peroxidation—a marker of reactive oxygen species induced stress, in which peroxidized lipids shift fluorescent emission peak from red (∼590nm) to green (∼510nm)^30^, together with microglial (Iba1) and astrocytic (GFAP) markers. Intriguingly, in PS19/E4 mice, peroxidized lipids (green) were found predominantly in Iba1^+^ microglia, while non-peroxidized lipids (red) localized primarily to GFAP^+^ astrocytes and, to a lesser extent, Iba1^+^ microglia (Fig. 1j-l). These findings indicate heightened lipid peroxidation specifically within microglia.

To further determine the subcellular localization of the BODIPY^+^ neutral lipids, we co-stained sections with markers of early endosomes (Rab5), late endosomes (Rab7), and lysosomes (LAMP1) (Extended Data Fig. 3h-j). We did not observe clear co-localization with Rab5 or Rab7 (Extended Data Fig. 3h,i), but ∼75% of BODIPY signal overlapped with LAMP1 in microglia (Extended Data Fig. 3j,k), indicating the majority of these neutral lipids reside within lysosomes. We speculate that the neutral lipids not co-localized with Rab5, Rab7, or LAMP1 in microglia may be cytosolic, potentially reflecting impaired phagocytic and lysosomal processing ability of APOE4 microglia^12,31,32^.

Taken together, these data show that many neutral lipids accumulated in PS19/E4 mice is localized to glial cells in the DG and exhibit spatial patterns distinct from LDs. The strong positive correlations between neutral lipid accumulation with gliosis, along with its inverse relationship with neuronal integrity, suggest that neutral lipids may serve as sensitive markers or even mediators of neurodegeneration in this model. The predominance of lipid peroxidation within microglia further suggest that microglia are key targets of oxidative stress-driven lipid dysregulation. These findings underscore a potential mechanistic link between neuronal APOE4, lipid dysregulation, and glial reactivity in vulnerable hippocampal circuits.

### Neuronal APOE4 is necessary and sufficient to induce damaging lipid accumulation

Cell-type-specific expression of APOE4 has become an important focus in AD research^23,32^, with neuronal APOE4 increasingly implicated in degenerative phenotypes^23,27,33–37^. Our lab previously generated a neuron-specific APOE4 deletion model, PS19/E4/Synapsin-1-Cre (PS19/E4/Syn1-Cre) mice, which exhibited significant reductions in tau pathology, gliosis, and neurodegeneration^23^. Using the same PS19/E4/Syn1-Cre mouse model, we found that neuronal APOE4 removal also markedly reduced both neuronal and microglial LDs (Fig. 2a-c). We also observed decreased neutral lipid accumulation in the hippocampal hilus of PS19/E4/Syn1-Cre compared to PS19/E4 mice (Fig. 2d,e). In PS19/E4/Syn1-Cre mice, neither microglial nor neuronal LDs nor neutral lipids showed significant correlations with other pathological endpoints (Extended Data Fig. 4a-l), except for a correlation between neuronal LDs and NeuN^+^ area coverage (Extended Data Fig. 4k). Overall, these findings demonstrate that neuronal APOE4 expression is necessary to drive the lipid accumulation and its associated pathological features in this tauopathy mouse model.

**Fig. 2.**
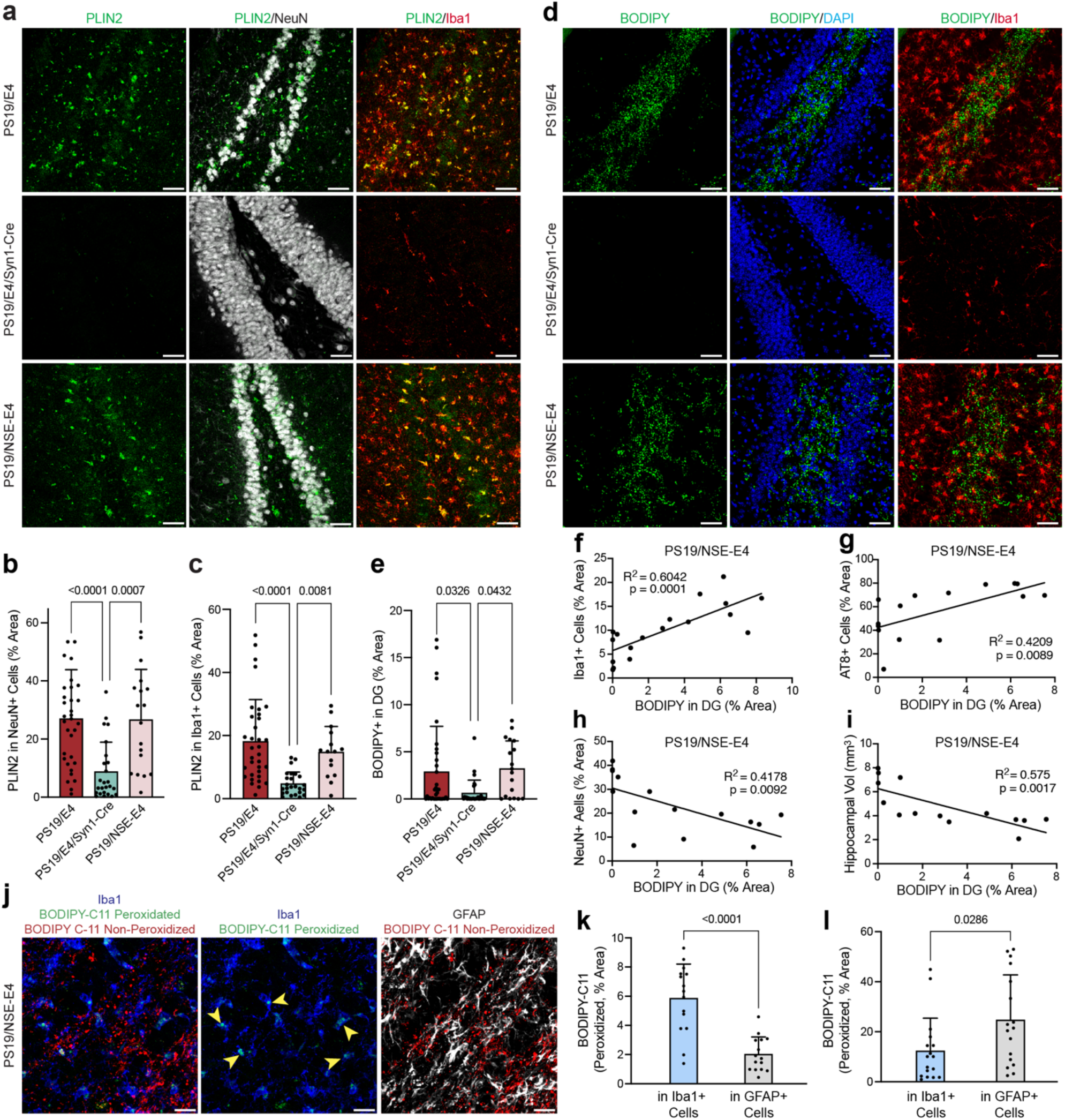
| Neuronal APOE4 is both necessary and sufficient to induce lipid accumulation in the hippocampal dentate gyrus (DG) in PS19 tauopathy mice. **a,** Representative immunofluorescent images of PLIN2^+^ lipid droplets (green), NeuN^+^ neurons (white), and Iba1^+^ microglia (red) in the hippocampal DG of 10-month-old PS19/E4, PS19/E4/Syn1-Cre, and PS19/NSE-E4 mice. Scale bars, 40μm. **b,** Quantification of percent PLIN2^+^ area coverage in NeuN^+^ neurons. **c,** Quantification of percent PLIN2^+^ area coverage in Iba1^+^ microglia. **d,** Representative immunofluorescent images of BODIPY^+^ neutral lipids (green), DAPI^+^ nuclei (white), and Iba1^+^ microglia (red) in the hippocampal DG of 10-month-old PS19/E4, PS19/E4/Syn1-Cre, and PS19/NSE-E4 mice. Scale bars, 40μm. **e,** Quantification of percent BODIPY^+^ area coverage in the hippocampal DG mask. Quantified data in **b-e** represented as mean ± s.e.m., unpaired two-sided t-test. **f-i,** Correlations between BODIPY^+^ % area in DG and Iba1^+^ % area (**f**), AT8^+^ % area (**g**), NeuN^+^ % area (**h**), and hippocampal volume (**i**) in PS19/NSE-E4 mice. **j,** Representative immunofluorescent images of BODIPY-C11^+^ peroxidized lipids (green), BODIPY-C11^+^ non-peroxidized lipids (red), Iba1^+^ microglia (blue), and GFAP^+^ astrocytes (white) in PS19/E4 mice. Scale bars, 20μm. **k,** Quantifications of BODIPY-C11^+^ peroxidized lipids in Iba1^+^ microglia and GFAP^+^ astrocytes in PS19/NSE-E4 mice. **l,** Quantifications of BODIPY-C11^+^ non-peroxidized lipids in Iba1^+^ microglia and GFAP^+^ astrocytes in PS19/NSE-E4 mice. Quantified data in **k** and **l** are represented as mean ± s.e.m., one-way analysis of variance (ANOVA) with Tukey’s post hoc multiple comparisons test. For all experiments, PS19/E4, *n*= 35; PS19/E3 *n*= 42; PS19/E4/Syn1-Cre *n*= 29; and PS19/NSE-E4 *n*= 18. Pearson’s correlation analysis (two-sided) used for **f-i**.

To determine whether neuronal APOE4 alone is sufficient to cause this lipid-accumulating phenotype, we used a previously reported mouse line expressing APOE4 under the neuron-specific enolase (NSE) promoter^38,39^. These NSE-E4 mice were cross-bred with an ApoE knockout background to ensure neuron-specific APOE4 expression, then further crossed with PS19 mice to generate PS19/NSE-E4 mice expressing APOE4 exclusively in neurons^27^. PS19/NSE-E4 mice exhibited lipid accumulation phenotypes similar to PS19/E4 mice, including significant LD buildup in neurons and microglia, as well as neutral lipid accumulation in microglia and astrocytes in the DG hilus (Fig. 2a-e). In PS19/NSE-E4 mice, neutral lipid accumulation positively correlated with both microgliosis and p-tau accumulation (Fig. 2f, g) and negatively correlated with NeuN^+^ percent area and hippocampal volume (Fig. 2h,i), mirroring the relationships observed in PS19/E4 mice. However, neuronal and microglial LDs did not correlate with most pathological measures (Extended Data Fig. 4m-t), except for PLIN2^+^/Iba1^+^ microglial LDs and NeuN^+^ percent area (Extended Data Fig. 4o). Similar to PS19/E4 mice, accumulated microglial lipids in PS19/NSE-E4 mice included both peroxidized and non-peroxidized lipids, whereas astrocytes primarily contained non-peroxidized lipids (Fig. 2j-l). These results indicate that neuron-specific APOE4 expression fully recapitulates the lipid accumulating phenotypes observed with global APOE4 expression. Together, we show that neuronal APOE4 alone is sufficient to induce the lipid accumulating phenotypes observed in the tauopathy mice with global APOE4 expression.

### Neuronal APOE4 induces lipid-laden oligodendrocytes that feed microglial lipid accumulation

Notably, neutral lipid staining revealed that not all BODIPY^+^ neutral lipids were colocalized with microglia or astrocytes (Fig. 2d,j), suggesting that additional cell types may contribute to the observed lipid pool. Given that recent studies increasingly implicate oligodendrocyte dysfunction^23,24,25^, myelin instability^25^, and altered oligodendrocytic lipid processing^25,26^ in both aging and APOE4 contexts, we asked whether oligodendrocytes might serve as an additional source of the neutral lipids in PS19/E4 and PS19/NSE-E4 hippocampi.

To address this question, we stained hippocampal sections from PS19/E4, PS19/E3, PS19/E4/Syn1-Cre, and PS19/NSE-E4 mice for Olig2 together with BODIPY. Interestingly and importantly, both global (PS19/E4) and neuron-specific APOE4 expression (PS19/NSE-E4) produced marked increases in BODIPY^+^ lipid accumulation within Olig2^+^ oligodendrocytes compared with PS19/E3 and PS19/E4/Syn1-Cre mice (Fig. 3a,b).

**Fig. 3.**
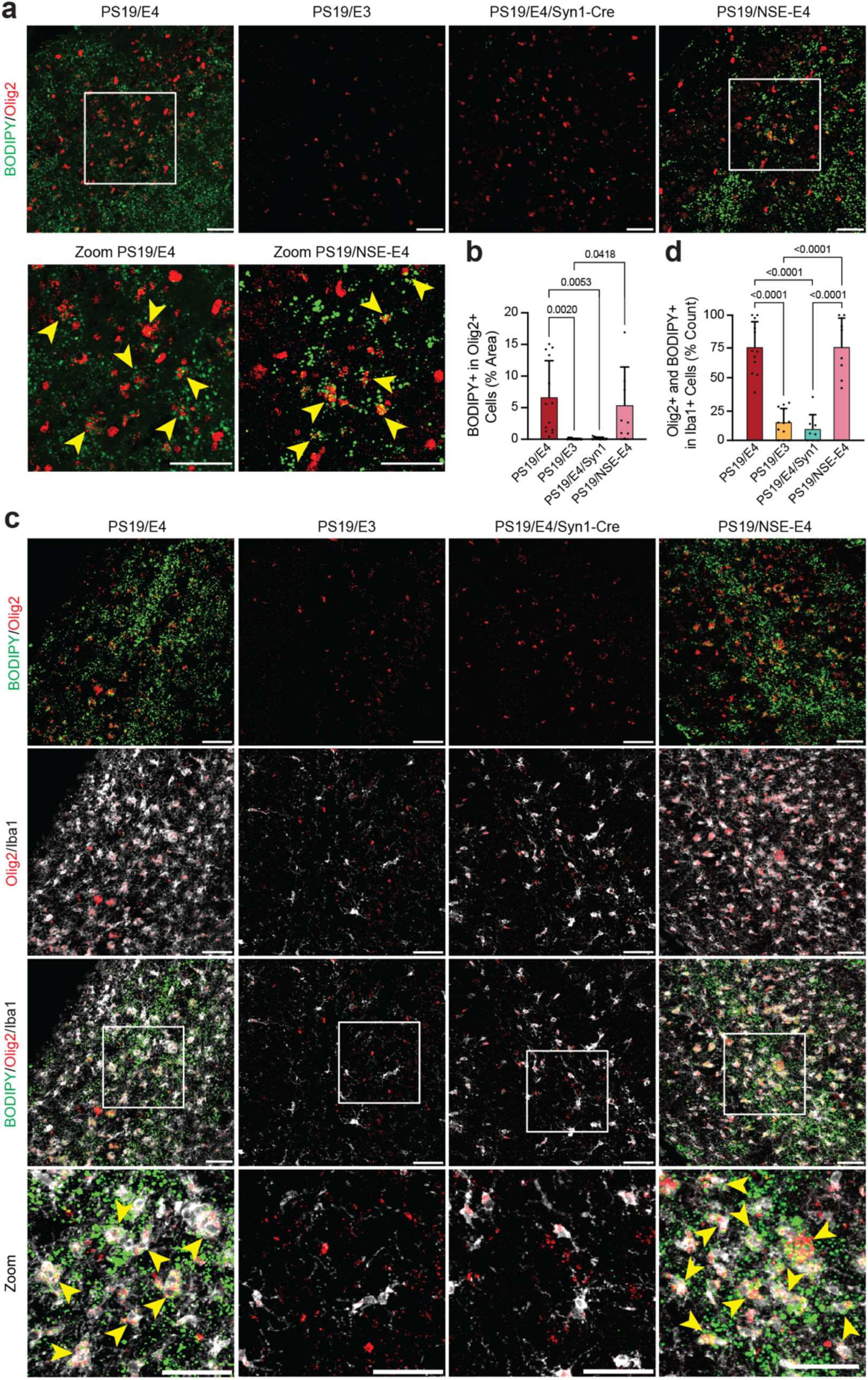
| Neuronal APOE4 acts through oligodendrocytes to influence microglial lipid accumulation. **a,** Representative immunofluorescent images of Olig2^+^ (red) oligodendrocytes co-localizing with BODIPY^+^ neutral lipids (green) in the hippocampal DG of 10-month-old PS19/E4, PS19/E4/Syn1-Cre, and PS19/NSE-E4 mice. Yellow arrows in zoomed in images represent co-localization of neutral lipids inside Olig2^+^ oligodendrocytes. Yellow arrows in zoomed-in images represent Olig2^+^ oligodendrocytes containing BODIPY^+^ neutral lipids. Scale bars, 40μm. **b,** Quantification of percent BODIPY^+^ area coverage in Olig2^+^ oligodendrocytes. (PS19/E4, *n*= 13; PS19/E3 *n*= 12; PS19/E4/Syn1-Cre *n*= 5, and PS19/NSE-E4 *n*= 7). **c,** Representative immunofluorescent images of BODIPY^+^ neutral lipids (green), Olig2^+^ oligodendrocytes (red), and Iba1^+^ microglia (white) in the hippocampal DG of 10-month-old PS19/E4, PS19/E4/Syn1-Cre, and PS19/NSE-E4 mice. Yellow arrows in zoomed-in images represent Iba1^+^ microglia containing both Olig2^+^ oligodendrocytes and BODIPY^+^ neutral lipids. Scale bars, 40μm. **d,** Quantification of percent of Iba1^+^ microglia that contain both Olig2^+^ oligodendrocytes and BODIPY^+^ neutral lipids, expressed as a percentage of total Iba1^+^ microglia. (PS19/E4 *n*= 12; PS19/E3 *n*= 12; PS19/E4/Syn1-Cre *n*= 8, and PS19/NSE-E4 *n*= 8). Quantified data in **b,d** are represented as mean ± s.e.m., one-way analysis of variance (ANOVA) with Tukey’s post hoc multiple comparisons test.

We next examined the morphological relationship between the lipid-laden oligodendrocytes and microglia by co-staining neutral lipids (BODIPY), oligodendrocytes (Olig2), and microglia (Iba1). In PS19/E4 and PS19/NSE-E4 mice, ∼75% of microglia were positive for both Olig2 and BODIPY, whereas only ∼10–20% of microglia stained positive for both markers in PS19/E3 and PS19/E4/Syn1-Cre mice (Fig. 3c,d). This pattern suggests that microglia likely engulf lipid-laden oligodendrocytes when neuronal APOE4 is expressed. Overall, these findings indicate that neuronal APOE4 not only drives lipid accumulation within oligodendrocytes but also promote their uptake by microglia, positioning oligodendrocytes as an additional neuronal APOE4-dependent source of lipid cargo delivered to microglia.

### Neuronal APOE4 orchestrates lipid accumulation via contact-dependent neuron-oligodendrocyte-microglia interaction

To investigate the cellular mechanisms by which neuronal APOE4 drives lipid accumulation, we performed co-culture experiments with primary neurons from PS19/E4 and PS19/E4/Syn1-Cre pups (PE4-N, expressing APOE4; and PE4/Syn1-N, lacking APOE) at postnatal day 0 (P0), and primary microglia from PS19/E4 (PE4-MG) pups at postnatal day 3 (P3). The cultured primary neurons were TUJ1^+^ with no detectable GFAP^+^ astrocytes or Iba1^+^ microglia (Extended Data Fig. 5a), while the primary microglia expressed canonical markers, including P2RY12, CD68, and Iba1 (Extended Data Fig. 5b).

First, we asked whether APOE4^+^ neurons transfer lipids to microglia through secreted factors by performing a medium transfer assay. Conditioned media from PE4-N (APOE4^+^) and PE4/Syn1-N (APOE4-null) was mixed 1:1 with microglia complete media and applied to PS19/E4 microglia for 72 hours. Immunocytochemistry (ICC) for PLIN2^+^ LDs and BODIPY^+^ neutral lipids revealed no differences between conditions (Extended Data Fig. 5c,d and Fig. 4a,b). In fact, lipid levels were low overall, indicating that microglial APOE4 does not account for the lipid accumulating phenotype in this *in vitro* system.

**Fig. 4.**
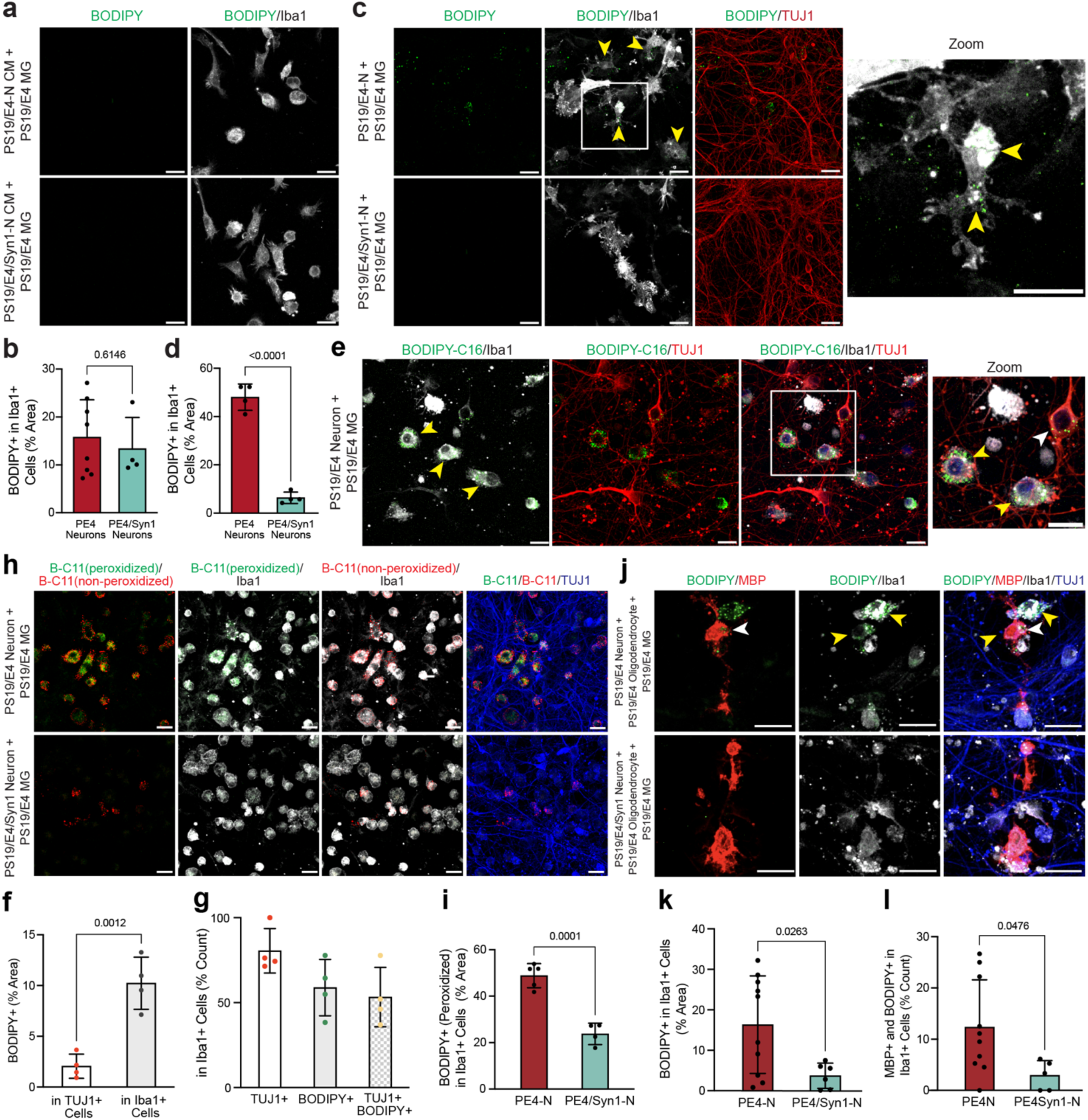
| Neuronal APOE4 promotes microglial lipid accumulation via contact-dependent neuron-oligodendrocyte-microglia interaction. **a,** Representative immunofluorescent images of BODIPY^+^ neutral lipids (green) and Iba1^+^ microglia (white) from PS19/E4 mice, treated with conditioned media (CM) from neurons of PS19/E4 or PS19/E4/Syn1-Cre mice for 72 hours. **b,** Quantification of percent BODIPY^+^ area coverage in Iba1^+^ microglia (MG) from PS19/E4 mice (PS19/E4 neuron CM, *n*= 8; PS19/E4/Syn1-Cre neuron CM *n*= 4). **c,** Representative immunofluorescent images of BODIPY^+^ neutral lipids (green), Iba1^+^ microglia (white), and TUJ1^+^ neurons (red) of PS19/E4 microglia co-cultured with PS19/E4 or PS19/E4/Syn1-Cre neurons for 72 hours. Arrows point to BODIPY in Iba1^+^ microglia. **d,** Quantification of percent BODIPY^+^ area coverage in Iba1^+^ microglia (MG) from PS19/E4 mice (PS19/E4 neuron, *n*= 4; PS19/E4/Syn1-Cre neuron, *n*= 4). **e,** Representative immunofluorescent images of BODIPY-C16^+^ lipids (green), Iba1^+^ microglia (white), and TUJ1^+^ neurons (red) from the lipid transfer assay with PS19/E4 microglia and PS19/E4 neurons. Yellow arrows point towards BODIPY-C16-lipids transferred from neurons to microglia and white arrow points toward BODIPY-C16-lipids remaining in neurons. (*n*= 4). **f,** Quantification of BODIPY-C16-lipids in TUJ1^+^ neurons (checkered) and Iba1^+^ microglia (stripe). (PS19/E4 neuron, *n*= 4; PS19/E4/Syn1-Cre neuron, *n*= 4). **g,** Quantification of count represented in percent of Iba1^+^ microglia containing TUJ1^+^ (red dots), BODIPY^+^ (green dots), and TUJ1^+^and BODIPY^+^ (yellow dots) signals. (PS19/E4, *n*= 4). **h,** Representative immunofluorescent images of peroxidized BODIPY-C11-lipids (B-C11, green), non-peroxidized BODIPY-C11-lipids (B-C11, red), Iba1^+^ microglia (white), and TUJ1^+^ neurons (blue) in mixed neuron-microglia co-culture. **i,** Quantifications of BODIPY-C11^+^ peroxidized lipids in Iba1^+^ microglia. (PS19/E4 neuron, *n*= 5; PS19/E4/Syn1-Cre neuron, *n*= 4). (PS19/E4 neuron, *n*= 5; PS19/E4/Syn1-Cre neuron, *n*= 4). **j,** Representative immunofluorescent images of BODIPY^+^ neutral lipids (green), MBP+ oligodendrocytes (red), Iba1^+^ microglia (white) and TUJ1^+^ neurons (blue) in mixed culture of PS19/E4 microglia, PS19/E4 oligodendrocytes, and either PS19/E4 or PS19/E4/Syn1-Cre neurons. Scale bars, 10μm. **k**, Quantification of percent BODIPY^+^ area coverage in Iba1^+^ microglia in the mixed culture of PS19/E4 microglia, PS19/E4 oligodendrocytes, and either PS19/E4 or PS19/E4/Syn1-Cre neurons. (PS19/E4 neuron, *n*= 10; PS19/E4/Syn1-Cre neuron, *n*= 6). **l,** Quantification of count represented in percent of Iba1^+^ microglia containing both MBP^+^ oligodendrocytes and BODIPY^+^ neutral lipids, expressed as a percentage of total Iba1^+^ microglia. (PS19/E4-Neuron Condition, *n*= 10; PS19/E4/Syn1-Cre Neuron Condition *n*= 5). Quantified data in **b,d,f,g,i,k,l** represented as mean ± s.e.m., unpaired two-sided t-test. In **a,c,e,h,j** all scale bars, 10μm.

We next tested whether physical contact between neurons and microglia is required for microglial lipid accumulating phenotype. Isolated PS19/E4 microglia were added onto PS19/E4 neurons (PE4-N + PE4-MG) or PS19/E4/Syn1-Cre neurons (PE4/Syn1-N + PE4-MG) and co-cultured for 72 hours. BODIPY staining showed a substantial increase in neutral lipid accumulation in microglia co-cultured with APOE4^+^ neurons, whereas lipid levels were significantly reduced when neurons lacked APOE4 (Fig. 4c,d). Neutral lipids primarily co-localized with Iba1^+^ microglia rather than TUJ1^+^ neurons, demonstrating that neuronal APOE4 drives microglial neutral lipid accumulation (Fig. 4c,d). Notably, PE4-N and PE4/Syn1-N neurons alone (at day 10, similar maturity as in co-culture) showed no differences in their intrinsic BODIPY^+^ or PLIN2^+^ lipid levels (Extended Data Fig. 5e-g). These results indicate that microglial lipid accumulation requires direct contact with neurons, consistent with microglia engulfing damaged neurons or synapses rather than receiving lipids through secretion.

To confirm that microglial lipid accumulation reflects uptake of neuronal lipids, we labeled neuronal lipids with BODIPY FL-C_16_ (BODIPY-C16), a fluorescent fatty acid analog that incorporates into newly synthesized lipids. Ten-day-old PS19/E4 neurons were pulsed with BODIPY-C16 for 1 hour and thoroughly washed to remove unincorporated dye, before adding PS19/E4 microglia for a 24-hour co-culture. Strikingly, the BODIPY-C16 labeled neuronal lipids were predominantly found in Iba1^+^ microglia, with only a low abundance in a subset of neurons (Fig. 4e,f), indicating that microglia were physically taking up lipids from neurons. Approximately half of the BODIPY-C16^+^ microglia also had TUJ1^+^ neuronal materials (Fig. 4e,g), indicating that microglia acquire neutral lipid through endocytosis and/or engulfment of neuronal components.

We next examined the effect of the neuronal APOE4 on lipid peroxidation in microglia. BODIPY-C11staining showed robust accumulation of both peroxidized (green) and non-peroxidized (red) lipids in microglia in the PE4-N + PE4-MG co-culture, whereas both signals were significantly reduced in PE4/Syn1-N + PE4-MG (Fig. 4h,i). Thus, neuronal APOE4 promotes both increased lipid uptake and increased lipid peroxidation within microglia.

Guided by our *in vivo* data, we next assessed the contribution of oligodendrocytes to microglial lipid accumulation. Using the mixed glial culture from the PS19/E4 pups, we isolated oligodendrocyte precursor cells (OPCs) using a previously described protocol^40^. After isolation, OPCs were co-cultured with PE4-N or PE4/Syn1-N and PE4-MG in a tri-culture system. After 72 hours, OPCs had matured into oligodendrocytes, as confirmed by extended processes in brightfield images (Extended Data Fig. 5h) and positive staining for myelin basic protein (MBP^+^) in the tri-culture (Fig. 4j), as well as Olig2^+^ in the monocultures (Extended Data Fig. 5i).

Strikingly, the PE4-N + PE4-MG + PS19/E4 Oligodendrocytes (PE4-O) tri-culture showed clear lipid accumulation in both oligodendrocytes and surrounding microglia, whereas the PE4/Syn1-N + PE4-MG + PE4-O tri-culture lacked lipid accumulation (Fig. 4j,k). Significantly more BODIPY^+^ microglia were observed surrounding oligodendrocytes and contained MBP^+^ material in tri-culture with APOE4^+^ neuron (Fig. 4j,k,l). The only variable between conditions was neuronal APOE4 expression, demonstrating that neuronal APOE4 is required for microglial uptake of oligodendrocyte-derived lipids.

Together, these *in vitro* findings from the neuron-oligodendrocyte-microglia tri-cultures show that neuronal APOE4 is both necessary and sufficient to drive microglial lipid accumulation through contact-dependent mechanism involving uptake of lipids from neurons and oligodendrocytes. It is also likely that neuronal APOE4 induces neuronal damage or degeneration, which in turn impairs oligodendrocyte health, thereby promoting microglial engulfment of lipid-rich material from both cell types.

### Microglial depletion reduces neutral lipid, but not LD, accumulation in the hippocampus

To further understand the role of microglia in regulating hippocampal lipid accumulation *in vivo*, we conducted a microglia depletion study in our PS19/E4 mice. Previously published data have shown that PLX3397 (a selective CSF1R/c-kit/FLT3 inhibitor) depletion of microglia reduces AD-related pathologies in PS19/E4 mice^36,41^. Following these studies, we treated PS19/E4 mice, starting at 7 months of age, with either control chow or PLX5622 chow, which is more selective and has higher brain penetration compared to PLX3397^42^, at 400ppm for 3 months. As expected, PLX5622-treated PS19/E4 (PS19/E4-PLX) mice exhibited ∼80% reduction in microglia without altering astrocytes levels (Extended Data Fig. 6a-d). Curiously, we found that, while microglial reduction had no effect on neuronal LD accumulation (Extended Data Fig. 6e,f), the PLX treatment did significantly decrease the amount of neutral lipid accumulation in the DG compared to PS19/E4-control mice (Extended Data Fig. 6g,h).

Together, these results indicate that neuronal LD formation occurs independently of microglia, whereas microglia are required for the deposition of neutral lipids—potentially reflecting a more disordered, disease-associated lipid state compared to the organized compartmentalization provided by LDs. These findings highlight a critical role for microglia in shaping the lipid landscape of the APOE4 brain: while neurons autonomously form LDs, microglia appear necessary to generate or amplify neutral lipids, suggesting that they function both as responders to and propagators of lipid-related pathology in a neuronal APOE4–dependent manner.

### Neuronal APOE4 drives neuronal vulnerability associated with lipid pathology

To investigate the cell-type-specific roles of APOE4 on the lipid accumulating phenotype at the transcriptomic level, we performed single-nucleus RNA-sequencing (snRNA-seq) on hippocampal tissues from 10-month-old mice of four genotypes: PS19/E4, PS19/E3, PS19/E4/Syn1-Cre, and PS19/NSE-E4. The dataset contained 133,936 nuclei covering 26,546 genes after normalization and quality control (Extended Data Fig. 7a,b). Clustering by the Louvain algorithm and visualization by Uniform Manifold Approximation and Projection (UMAP) revealed 36 distinct cell clusters (Fig. 5a). Based on marker gene expression, these nuclei were assigned to excitatory (Ex.) neuron DG clusters (1, 11, 20, 29, 32), Ex. neuron CA1 clusters (4, 6, 9, 26, 28, 33, 34, 35), Ex. neuron CA2/3 clusters (8, 17, 27, 30), subiculum neuron clusters (5, 13, 18, 19, 21, 22), inhibitory neuron clusters (12, 31), oligodendrocyte clusters (2, 3, 7, 23), oligodendrocyte progenitor cell (OPC) clusters (16, 36), microglia clusters (10, 14), astrocyte clusters (15, 24), and a choroid plexus cluster (25) (Fig. 5a).

**Fig. 5.**
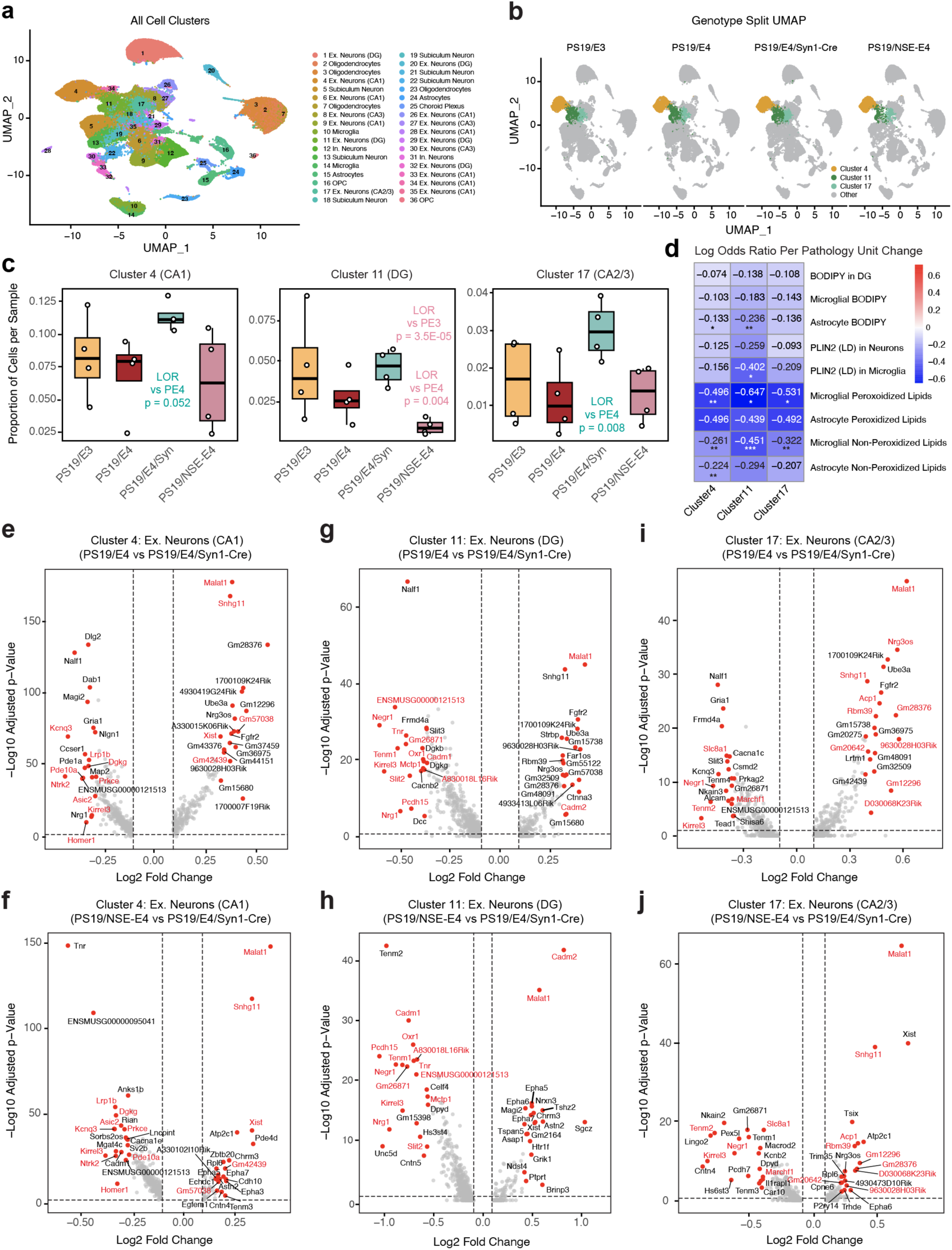
| Neuronal APOE4 diminishes protective excitatory neuron clusters. **a,** UMAP plot of all 36 distinct cell clusters in the hippocampi of 10-month-old PS19/E4, PS19/E3, PS19/E4/Syn1-Cre, and PS19/NSE-E4 mice (*n*= 4 for each genotype). **b,** Genotype-split UMAP plot highlighting cells in excitatory neuron clusters 4, 11, and 17 for each genotype group. **c,** Box plots of proportion of cells from each sample in excitatory neuron clusters 4, 11, and 17. The lower, middle, and upper hinges of the box plot correspond to the 25^th^, 50^th^, and 75^th^ percentiles, respectively. The upper whisker of the box plot extends from the upper hinge to the largest value no further than 1.5 x IQR from the upper hinge. The lower whisker extends from the lower hinge to the smallest values at most 1.5 x IQR from the lower hinge. IQR, interquartile range or distance between 25^th^ and 75^th^ percentiles. Data beyond the end of the whiskers are the outlier points. The LORs are the mean ± s.e.m. estimates of LOR for these clusters, which represents the change in the log odds of cells per sample from PS19/E3, PS19/E4/Syn1-Cre, or PS19/NSE-E4 mice belonging to the respective clusters compared to the log odds of cells per sample from PS19-E4 mice. PS19/E3 is represented in yellow, PS19/E4 is represented in red, PS19/E4/Syn1-Cre is represented in green, and PS19/NSE-E4 is represented in pink. LOR significant differences from either PS19/E4 or PS19/E3 are highlighted in corresponding color of the genotype (e.g. if PS19/NSE-E4 is significantly different, text is colored pink). **d,** Heat map plot of log odds ratio (LOR) per unit change in each pathological measurement for clusters 4, 11, and 17. The LOR represents the mean estimate of the change in the log odds of cells per sample from a given animal model, corresponding to a unit change in a given histopathological parameter. Associations with pathologies are colored (negative associations, blue; positive associations, red). P values in **c** are from fits to a GLMM_AM, and p values in **d** are from fits to a GLMM_histopathology; the associated tests are two-sided. All error bars represent s.e.m. * p < 0.05, ** p <0.01, *** p <0.001. **e,g,i,** Volcano plot for the top 40 (20 up and 20 down) differentially expressed genes (DEGs) of excitatory neuron cluster 4 (**e**), 11 (**g**), and 17 (**i**) for PS19/E4 versus PS19/E4/Syn1-Cre. **f,h,j,** Volcano plot for the top 20 DEGs of excitatory neuron cluster 4 (**f**), 11 (**h**), and 17 (**j**) for PS19/NSE-E4 versus PS19/E4/Syn1-Cre.

To evaluate the effect of neuronal APOE4, we used logs odds ratio (LOR) estimates from generalized linear mixed-effects models associated with animal models (GLMM_AM) to identify cell clusters that were altered in PS19/E4 versus PS19/E3, PS19/E4/Syn1-Cre, and PS19/NSE-E4. This analysis revealed that Ex. neuron clusters 4 (CA1) and 17 (CA2/3) had significantly higher odds of containing cells from PS19/E4/Syn1-Cre mice compared to PS19/E4 mice (Fig. 5b,c), suggesting their vulnerability to neuronal APOE4 toxicity. Importantly, Ex. Neuron clusters 4 and 17 correlated negatively with most neutral lipid measurements—and a significant negative association with microglial peroxidized lipids (Fig. 5d and Extended Data Fig. 8)— as determined with a GLMM associated with lipid pathology (GLMM_LP). This suggests these are lipid phenotype-protective neuron clusters that are vulnerable to neuronal APOE4 toxicity. Differentially expressed gene (DEG) analyses of PS19/E4 or PS19/NSE-E4 versus PS19/E4/Syn1-Cre mice revealed that cells in neuron clusters 4 and 17 significantly upregulated long-noncoding RNA (lncRNA) genes (*Malat1* and *Snhg11,* and *Xist* for cluster 4) and signal transduction (*Acp1* for cluster 17). In contrast, these neurons downregulated genes associated with synaptic signaling and plasticity (*Homer1 and Ntrk2*) and membrane potential modulation (*Kcnq3*) in cluster 4, and synaptic adhesion (*Kirrel3*) and neurite growth (*Negr1*) in cluster 17 (Fig. 5e,f,i,j), indicating that these transcriptional programs are particularly sensitive to neuronal APOE4 effects.

Ex. neuron cluster 11 (DG) showed significantly lower odds of containing cells from PS19/NSE-E4 compared to PS19/E3 and PS19/E4 mice and trended toward higher odds in PS19/E4/Syn1-Cre mice (Fig. 5b,c). Strikingly, this DG neuron cluster had the strongest negative correlation with many of the neutral lipid measurements (Fig. 5d), indicating that it is the strongest lipid phenotype-protective neuron cluster that is particularly vulnerable to neuronal APOE4 toxicity. Given the robust lipid phenotype observed within the DG in our *in vivo* models, these results were not surprising. Comparisons of DEGs in cluster 11 showed downregulation in structural and adhesive architecture genes (*Tenm1, Negr1,* and *Tnr*) and guidance and signaling refinement genes (*Nrg1* and *Slit2*), and upregulation in genes involved in synaptic regulation and transcriptional modulation (*Malat1* and *Cadm2*) (Fig. 5g,h). Strikingly, many DEGs were identical in PS19/E4 versus PS19/E4/Syn1-Cre and PS19/NSE-E4 versus PS19/E4/Syn1-Cre comparisons (Fig. 5e-j), suggesting that these effects are driven by neuronal APOE4.

Together, these findings suggest that neuronal APOE4 removal preserves or protects against the loss of clusters 4, 11, and 17, whereas neuronal APOE4 expression selectively depletes DG neurons. Collectively, clusters 4, 11, and 17 represent neuronal populations particularly vulnerable to neuronal APOE4 effects. Notably, these cell clusters have transcriptional programs converging on synaptic structure, connectivity, and plasticity, highlighting cellular processes that likely underlie their susceptibility to neuronal APOE4.

### Neuronal APOE4 drives disease associated oligodendrocyte and microglia contributing to lipid pathology

In addition to the vulnerable excitatory neuron populations, GLMM_LP analysis also revealed oligodendrocyte and microglia clusters that were positively associated with lipid pathologies (Fig. 6a,c and Extended Data Fig. 8). Oligodendrocyte cluster 7 showed a significant enrichment in PS19/NSE-E4 relative to PS19/E3 mice (Fig. 6a,b), resembling previously identified neuronal APOE4-induced disease-associated oligodendrocyte (nE4-DAO)^23,24,27^. Strikingly, oligodendrocyte cluster 7 showed strong positive associations with most lipid pathologies (Fig. 6c), highlighting a disease-associated state. DEG analyses of PS19/E4 or PS19/NSE-E4 versus PS19/E4/Syn1-Cre mice in cluster 7 showed similar upregulated and downregulated genes (Fig. 6d,e), suggesting that these gene expression changes are driven by neuronal APOE4. The upregulated genes included stress-induced lncRNA *Neat1* and RNA/protein processing genes (*Fmn1* and *Fbxw15*), and the downregulated genes included cell-cell adhesion (*Lsamp* and *Adgrb3*) and myelin-related signaling (*Lingo2, Epha6,* and *Cntnap2*) genes (Fig. 6d,e). This transcriptional profile reflects a shift to a stress-responsive and maladaptive state, likely in response to neuronal APOE4-induced damage of the APOE4 vulnerable neuronal clusters (such as clusters 4, 11, and 17).

**Fig. 6.**
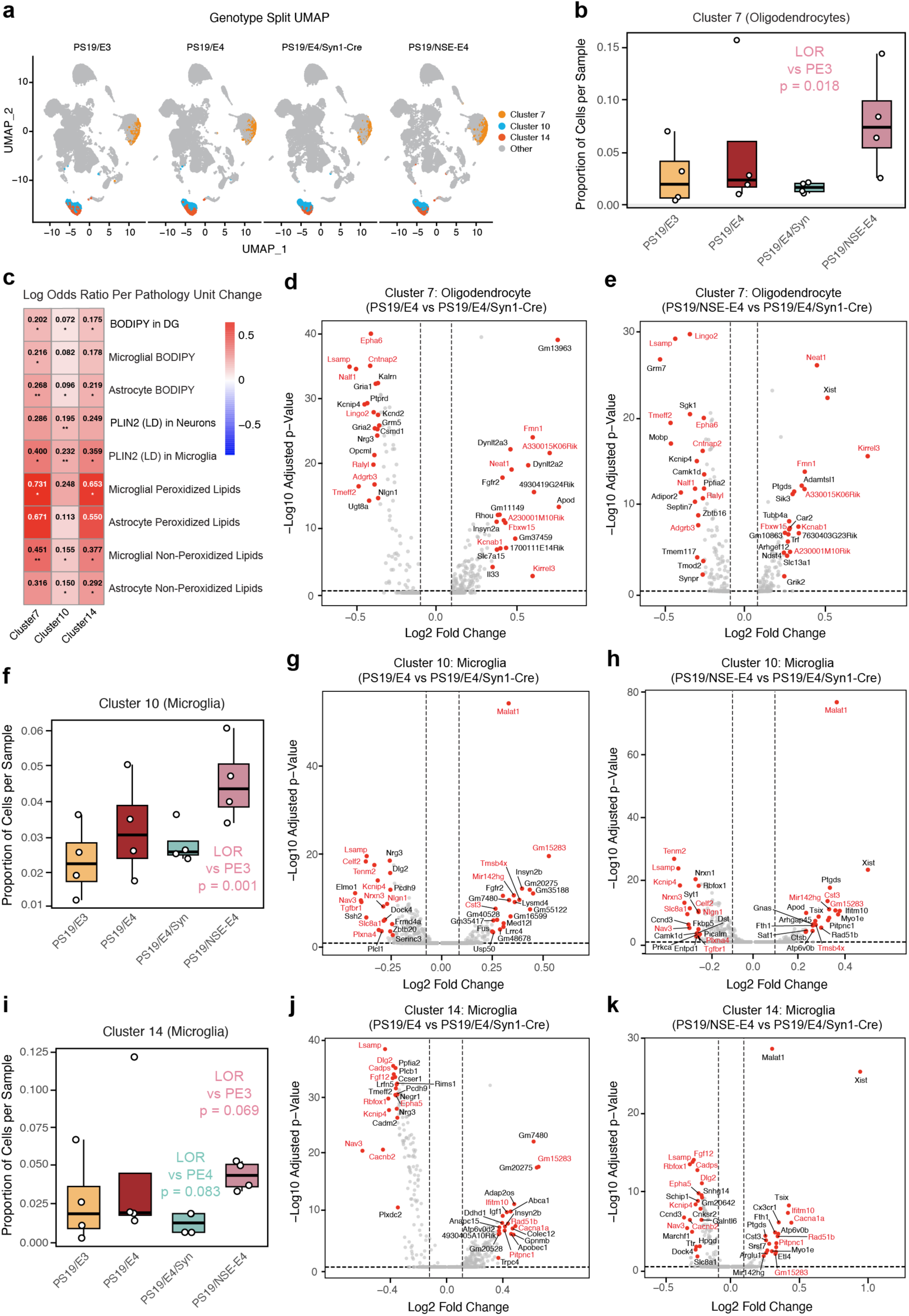
| Neuronal APOE4 increases lipid related disease-associated oligodendrocyte and microglial clusters. **a,** Genotype-split UMAP plot highlighting cells in oligodendrocyte cluster 7 and microglial clusters 10 and 14 for each genotype group. **b,** Box plot of proportion of cells from each sample in cluster 7. **c,** Heat map plot of log odds ratio (LOR) per unit change in each pathological measurement for clusters 7, 10, and 14. The LOR represents the mean estimate of the change in the log odds of cells per sample from a given animal model, corresponding to a unit change in a given histopathological parameter. Associations with pathologies are colored (negative associations, blue; positive associations, red). **d,e,** Volcano plot for the top 40 (20 up and 20 down) differentially expressed genes (DEGs) of oligodendrocyte cluster 7 for PS19/E4 versus PS19/E4/Syn1-Cre (**d**) and for PS19/NSE-E4 vs PS19/E4/Syn1-Cre (**e**). **f,** Box plot of proportion of cells from each sample in cluster 10. **g,h,** Volcano plot for the top 40 (20 up and 20 down) DEGs of microglia cluster 10 for PS19/E4 versus PS19/E4/Syn1-Cre (**g**) and for PS19/NSE-E4 vs PS19/E4/Syn1-Cre (**h**). **i,** Box plot of proportion of cells from each sample in cluster 14. **j,k,** Volcano plot for the top 40 (20 up and 20 down) DEGs of microglia cluster 14 (**j**) for PS19/E4 versus PS19/E4/Syn1-Cre and for PS19/NSE-E4 vs PS19/E4/Syn1-Cre (**k**). For **b,f,i,** the lower, middle, and upper hinges of the box plot correspond to the 25^th^, 50^th^, and 75^th^ percentiles, respectively. The upper whisker of the box plot extends from the upper hinge to the largest value no further than 1.5 x IQR from the upper hinge. The lower whisker extends from the lower hinge to the smallest values at most 1.5 x IQR from the lower hinge. IQR, interquartile range or distance between 25^th^ and 75^th^ percentiles. Data beyond the end of the whiskers are the outlier points. The LORs are the mean ± s.e.m. estimates of LOR for these clusters, which represents the change in the log odds of cells per sample from PS19/E3, PS19/E4/Syn1-Cre, or PS19/NSE-E4 mice belonging to the respective clusters compared to the log odds of cells per sample from PS19-E4 mice. PS19/E3 is represented in yellow, PS19/E4 is represented in red, PS19/E4/Syn1-Cre is represented in green, and PS19/NSE-E4 is represented in pink. P values in **b,f,i** are from fits to a GLMM_AM, and p values in **c** are from fits to a GLMM_histopathology; the associated tests are two-sided. All error bars represent s.e.m. * p < 0.05, ** p <0.01, *** p <0.001.

Microglia clusters 10 and 14 had significantly increased odds of containing cells from PS19/NSE-E4 compared to PS19/E3 (Fig. 6f,i), suggesting that these microglial subpopulations are increased in response to neuronal APOE4-driven pathology. Strikingly, both microglial clusters 10 and 14 showed significant positive associations with most lipid pathologies (Fig. 6c), highlighting a disease-associated state. DEG analysis of PS19/E4 or PS19/NSE-E4 versus PS19/E4/Syn1-Cre mice revealed highly similar DEG profiles for both microglial clusters 10 and 14 (Fig. 6g,h,j,k), indicating that neuronal APOE4 alone is sufficient to elicit a microglial state comparable to that induced by global APOE4 expression. In cluster 10, both comparisons demonstrated coordinated downregulation of neuronal-interaction and ion-homeostasis–related transcripts (including *Lsamp, Slc8a1, Nrxn3,* and *Tenm2*) together with upregulation of stress-associated lncRNAs (*Mir142hg* and *Malat1*) (Fig. 6g,h), consistent with an early reactive microglial response. In contrast, both comparisons in cluster 14 exhibited shared downregulation of synaptic-support and RNA-processing genes (*Dlg2, Rbfox1, Cadps,* and *Cacnb2*) accompanied by upregulation of calcium-signaling and interferon/stress-response genes (*Cacna1a, Rad51b*, and *Ifitm10*) (Fig. 6j,k), hallmark features of a DAM-like, stress-responsive microglial state^43–45^.

Together, these findings indicate that neuronal APOE4 not only drives lipid-associated vulnerability in excitatory neurons but also induces stress-responsive, disease-associated states in both oligodendrocytes and microglia. Oligodendrocyte cluster 7 and microglial clusters 10 and 14 show transcriptional and lipid-pathology profiles consistent with maladaptive responses to neuronal APOE4, highlighting a coordinated glial reaction. These results underscore the broad, non-cell-autonomous impact of neuronal APOE4 on the cellular lipid landscape of the PS19 brain.

## Discussion

This study identifies neuronal APOE4 as a central conductor of lipid dysregulation that propagates through a neuron–oligodendrocyte-microglia interactive network to amplify tau-mediated neurodegeneration. In the PS19 tauopathy background, APOE4 drives robust accumulation of LDs and neutral lipids in the hippocampal dentate gyrus, with peroxidized lipids predominating in microglia. Further characterization showed that neuronal APOE4 is both necessary and sufficient for this phenotype: neuron-specific removal of APOE4 abrogates lipid accumulation and neuron-specific expression of APOE4 is sufficient to drive the disease phenotypes. Additional investigation reveals that microglial engulfment of damaged neurons and lipid-rich oligodendrocytes contributes to this lipid accumulating phenotype. snRNA-seq analysis further reveals that neuronal APOE4 reshapes cellular composition and state across the hippocampus—depleting protective excitatory neuron subclusters (clusters 4, 11, 17) while promoting lipid- and disease-associated oligodendrocyte (cluster 7) and microglia (clusters 10 and 14) populations.

The temporal dynamics of lipid accumulation suggests a stepwise pathogenic process. Neurons in PS19/E4 mice accumulate LDs linearly with age, reflecting gradual metabolic stress, whereas microglial lipid accumulation rises exponentially and coincide with gliosis onset^28^. This sequence indicates that neuronal stress precedes and drives microglial dysfunction, further adding to previous studies which highlight the importance of neuron-glia interactions for lipid dysregulation^46–49^. The engagement of oligodendrocytes as an additional lipid reservoir extends this model to a tri-cellular cascade, whereby neuronal APOE4 initiates multicellular-system lipid imbalance that propagates through interconnected neuronal, oligodendrocytic, and microglial compartments.

Our *in vitro* co-culture experiments further revealed that microglial lipid accumulation depends on physical contact with APOE4-expressing neurons rather than exposure to conditioned media, pointing to a phagocytosis (engulfment)-driven process of lipid transfer from neurons to microglia. The uptake of labeled neuronal lipids and the predominance of peroxidized lipids within microglia support a model in which neuronal APOE4 induces lipid transfer and oxidative stress, rendering neurons vulnerable and providing lipid-rich substrates for microglial engulfment and clearance. *In vitro* tri-culture experiments further support neuronal APOE4-damaged oligodendrocytes as an additional lipid source for microglial engulfment, aligning with the DAO signature and linking gray- and white-matter pathologies. This contact dependent mechanism was further validated *in vivo* where we observed microglia co-localizing with oligodendrocytes and neutral lipids, supporting a phagocytic/engulfment-dependent progression of lipid accumulation in microglia. Consistent with a central effector role for microglia, their depletion reduces neutral lipid burden without altering neuronal LDs, indicating that neuronal LD formation is microglia-independent whereas neutral lipid accumulation depends on microglial presence and processing.

snRNA-sequencing data supports a neuron-first model in which neuronal APOE4 reduces protective excitatory neuron populations (clusters 4, 11, and 17) while expanding disease-associated glial states linked to lipid pathology. The diminished neuronal clusters are enriched for synaptic adhesion, plasticity, and axon-guidance programs, emphasizing the selective vulnerability of highly interconnected neuronal circuits to APOE4-driven metabolic stress. In contrast, neuronal APOE4 induces a DAO cluster that is enriched in PS19/NSE-E4 mice and exhibits a transcriptional signature of the neuronal APOE4–associated DAO state^23,27^, including increased stress-response genes (*Neat1, Fmn1,* and *Fbxw15*) and reduced adhesion and myelin-related signaling (*Lsamp, Adgrb3, Lingo2, Epha6,* and *Cntnap2*). Oligodendrocytes are strongly associated with multiple lipid pathologies, indicating their contribution to APOE4-driven lipid dysfunction in AD.

Neuronal APOE4 also induces two distinct microglial activation states. Cluster 10 shows early stress reactivity characterized by downregulation of neuronal-interaction transcripts and upregulation of stress-associated lncRNAs, with neuronal APOE4 uniquely inducing lipid- and membrane-related genes (*Ptgds* and *Myo1e*). Cluster 14 represents a more DAM-like cluster marked by calcium signaling, interferon-stress responses, and loss of synaptic-support and RNA-processing genes. Together, these clusters delineate a shift from early activation to a lipid-engaged, stress-amplified microglial state driven specifically by neuronal APOE4.

Collectively, these findings show that neuronal APOE4 reprograms both oligodendrocytes and microglia into lipid-engaged, disease-associated states while depleting vulnerable neuronal populations. This neuron-initiated cascade disrupts glial homeostasis, amplifies lipid stress across cell types, and establishes a feed-forward circuit of glial dysfunction that accelerates APOE4-dependent neurodegeneration.

Our findings support a model in which neuronal APOE4 toxicity extends beyond direct neuronal dysfunction to orchestrate brain-wide glial collapse and lipid pathology (Extended Data Fig. 9). Neurons expressing APOE4 provide the initiating lipid stress, oligodendrocytes amplify this dysfunction by failing to maintain their integrity and providing lipid cargo, and microglia emerge as the final effector cell type that uptakes, accumulates, and oxidizes these lipids. The resulting lipid-laden, peroxidized microglia adopt transcriptional programs that are disease-associated (DAM), which may further drive inflammation and neurodegeneration. This model places neuronal APOE4 as an upstream and central orchestrator of multi-cellular lipid dysregulation: neuron-initiated, contact-dependent lipid transfer from neurons and oligodendrocytes to microglia reprograms microglia to disease-associated states, exacerbating neuroinflammation and contributing to neurodegeneration (Extended Data Fig. 9). These insights refine the cellular logic of APOE4 risk beyond global APOE-centric models^50–52^, positioning neuron-derived APOE4 as the upstream driver that organizes glial response, dysfunction, and lipid pathology. The observation that neuronal APOE4 removal simultaneously diminishes DAO and lipid-engaged, disease-associated microglia further highlights the causal leverage of the neuronal source of APOE4.

Our study also has some limitations. The mechanistic work is in a tauopathy mouse model with a hippocampal focus; validation in human APOE4 brain tissues and across regions will be an important next step. The subcellular localization and biochemical identity of the neutral lipid pools remain incompletely resolved and need further exploration. Finally, while microglial depletion implicates microglia in shaping neutral lipid landscapes, cell-type-specific perturbations of lipid processing genes will be required to separate causal roles in injury versus repair.

Nevertheless, our data suggest concrete therapeutic strategies. First, reducing the neuronal APOE4 driver (e.g., cell-type-specific ASOs/siRNA^53^, CRISPR base/prime editing^54^, or promoter-targeted repression^55^) to mitigate upstream lipid stress. Second, blocking neuron-to-microglia lipid transfer by modulating phagocytic pathways (e.g., complement, TREM2/APOE axis, integrin/adhesion cues) while preserving surveillance. Third, enhancing lipid efflux and clearance in glia via ABCA1/ABCG1 induction^18^, lysosomal biogenesis (TFEB/MITF), and LYST-dependent trafficking. Fourth, mitigating lipid peroxidation using lipid-targeted antioxidants (e.g., ferroptosis inhibitors, radical-trapping agents) and glutathione pathway support (xCT/SLC7A11). Finally, stabilizing oligodendrocyte function by promoting remyelination programs and correcting DAO-linked stress responses. By aligning interventions to combat the sequential process of neuronal APOE4 → neuronal lipid stress and damage → oligodendrocyte lipid accumulation and dysfunction → lipid-engaged/LDAM-like microglia, it may restore homeostatic neuron–oligodendrocyte-microglia interactions and ameliorate APOE4-driven neuroinflammation and neurodegeneration in AD and other neurodegenerative disorders.

## Methods

### Mice

Human LoxP-floxed APOE knock-in (E) mice with conditional deletion of the human APOE gene were generated as previously described. Briefly, homozygous E3 and E4 mice were crossed with Synapsin-1-Cre transgenic mice^23^ (B6.Cg-Tg(Syn1-Cre)671Jxm/J; The Jackson Laboratory, 003966). The resulting E/Cre mice were subsequently bred with tau-P301S (PS19) transgenic mice (B6;C3-Tg(Prnp-MAPT*P301S)PS19Vle/J; The Jackson Laboratory, 008169), which express the human 1N4R tau isoform with the P301S mutation under the PrP promoter, to generate PS19/E3 and PS19/E4 mice either with or without Syn1-Cre, as previously described^23^. Littermates lacking Syn1-Cre were used as PS19/E controls. To avoid germline recombination, only female Syn1-Cre mice were used for breeding. We also generated mice with APOE expressed under the neuron-specific enolase (NSE) promoter, and these mice were bred with an APOE knockout (KO) background mice to ensure APOE expression only in the neurons. These were then crossed with the PS19 mice to generate PS19/NSE-E4 mice^27,38,39^. We also bred the PS19 mice on the APOE KO background to generate PS19/EKO mice. All mice were maintained on a C57BL/6 background and housed in a pathogen-free barrier facility under standard conditions (12-hour light/dark cycle, 19–23 °C, 30–70% humidity). Mice were identified via ear punch under brief isoflurane anesthesia, and genotypes were confirmed by PCR analysis of tail clippings. No procedures were performed on animals beyond those described in this study. Both male and female mice were used for all experiments. All procedures were conducted in compliance with NIH guidelines and approved by the University of California, San Francisco Institutional Animal Care and Use Committee (protocol AN176773), under the oversight of the Laboratory Animal Resource Center at UCSF and the Gladstone Institutes.

PS19-E mice were analyzed at 10 months of age. Mice were anesthetized with intraperitoneal injections of avertin (Henry Schein) and transcardially perfused with 0.9% saline for 1 minute. Brains were dissected and processed as hemi-brains. Right hemispheres were drop-fixed in 4% paraformaldehyde (PFA, Electron Microscopy Sciences) for 48 hours, washed in 1× PBS (Corning) for 24 hours, and cryoprotected in 30% sucrose (Sigma) for 48 hours at 4 °C. Fixed hemispheres were sectioned coronally at 30 µm using a freeze-sliding microtome (Leica) and stored at −20 °C in cryoprotectant solution (30% ethylene glycol, 30% glycerol, and 40% 1× PBS). Left hemispheres were snap frozen on dry ice and stored at −80 °C for biochemical analyses or snRNA processing.

### PLX5622 administration

PLX5622 was provided by MedChemExpress, where the compound was incorporated into AIN-76A chow (Research Diets) at a concentration of 400ppm. PS19-fE4 mice were fed with either control AIN-76A or PLX5622 chow for 3 months, starting at 7 months until perfusions at 10 months of age. Brains were dissected and processed as hemi-brains as described above.

### In vivo immunohistochemistry (IHC) and BODIPY staining

Free floating brain sections were washed with 2× PBS being incubated in blocking solution (5% normal donkey serum (NDS) (017000121, Jackson Immuno), 0.2% Triton-X (Millipore Sigma) with Mouse-on-Mouse (M.O.M.) Blocking Buffer (MKB-2213, Vector Labs): one drop M.O.M IgG in 4 ml PBS in 1× PBS for 1 h at room temperature at room temperature. After MOM block, sections were incubated in primary antibody at 4 °C overnight after being diluted to optimal concentrations for the following proteins: anti-PLIN2 (AB52356, Abcam, 1:1,000); anti-Iba1(rb) (AB178846, Abcam, 1:1,000); anti-Iba1 (gp) (234308, Synaptic Systems, 1:500); anti-Iba1 (gt) (AB5076, Abcam, 1:1,000); anti-NeuN (ABN90, Milipore Sigma, 1:500); anti-GFAP (MAB3402, EDM Milipore, 1:800); anti-Olig2 (AF2418, R&D Systems, 1:100); and anti-MBP (AB7349, Abcam, 1:500). After primary antibody incubation, sections were washed in PBS and incubated at room temperature for 1 h in secondary antibodies: donkey anti-rabbit 405 (AB175651, Abcam, 1:1,000); donkey anti-guinea pig 405 (706-475-148, Jackson Immuno, 1:1,000); donkey anti-mouse 488 (A21202, Invitrogen, 1:1,000); donkey anti-rabbit 488 (A21206, Invitrogen, 1:1,000); donkey anti-mouse 594 (A21203, Invitrogen, 1:1,000); donkey anti-rabbit 594 (A21207, Invitrogen, 1:1,000); donkey anti-rat 594 (A21209, Invitrogen, 1:1,000); donkey anti-mouse 647 (A31571, Invitrogen, 1:1,000); donkey anti-rabbit 647 (A31573, Invitrogen, 1:1,000); and donkey anti-guinea pig 647 (706-65-148, Jackon Immuno, 1:1,000). Secondaries antibodies were co-incubated with DAPI (ThermoFisher, 1:20,000 dilution in PBS). For neutral lipid staining, BODIPY 493/503 (D3922, ThermoFisher, 1:1,000 dilution from 1mg/mL stock solution in DMSO) was added to secondary antibody incubation where the 488 channel was left empty for BODIPY imaging. For peroxidated lipid staining, BODIPY 581/591 C11 (BODIPY-C11) (Lipid Peroxidation Sensor) (D3861, Invitrogen, 1:1,000) was added to secondary antibody incubation where both the 488 and 594 channels were left empty for BODIPY imaging.

For successful neutral lipid and lipid droplet staining, detergent washes were avoided except for permeabilization with PBS-TX for the blocking step. After washing with PBS, sections were mounted onto microscope slides (Fisher Scientific), and coverslipped with ProLong Gold mounting medium (P36930, Vector Labs). Images were taken using an FV3000 confocal laser scanning microscope (Olympus) at ×20, ×40, or ×60 magnification depending on the stain. For each stain, all samples were imaged at the same fluorescent intensity.

For diaminobenzidine (DAB) staining, 30um-thick brain sections were washed three times with PBS-T, incubated in boiling citrate buffer (10mM sodium citrate; BP327-1, Fisher Bioreagents) for 5 min, washed two times with PBS-T, incubated in endogenous peroxidase block (3% H2O2; H1009, Millipore Sigma and 10% methanol; A412SK-4, Fisher Scientific in PBS) for 15 min, washed two more times with PBS-T, washed one time with PBS-Tx (PBS + 0.2% TritonX; T8787, Millipore Sigma), followed by blocking with 10% NDS made in PBS-Tx for 1 hr. Sections were then incubated in avidin and biotin blocks (3 drops of each; SP-2001, Vector Laboratories) for 15 min and MOM Blocking buffer (one drop MOM per 4 ml of PBS-T) for 1 hr. Sections were incubated overnight at 4°C with an anti-AT8 primary antibody (MN1020, Invitrogen, 1:100) in 3% NDS made in PBS-T. The following day, sections were washed three times with PBS-T and incubated for 1 hr at room temperature with biotinylated donkey anti-mouse secondary (715-065-150, Jackon ImmunoResearch, 1:2,000) in 3% NDS in PBS-T. Next, the sections were rinsed two times with PBS-T, once with PBS, and incubated with ABC buffer (2 drops A and B per 5ml PBS; PK-6100, Vector Laboratories) for 1 hr. The ABC buffer was made and allowed to incubate 15-30 min before use. Next, the sections were rinsed twice times with PBS and once with water before a 1 min and 30 sec incubation in DAB buffer (2 drops buffer stock solution, 4 drops DAB, and 2 drops H_2_O_2_ in 5ml Milli-Q water; SK-4100, Vector Laboratories). Sections were then immediately washed three times with Milli-Q water and then washed with PBS and mounted onto microscope slides. Slides were allowed to dry overnight and then submerged in two washes of xylene (HC7001GAL, Fisher Scientific). Slides were coverslipped with DPX mounting medium (06522, Millipore Sigma) and imaged with an Aperio VERSA slide scanning microscope (Leica).

### Volumetric analysis & neuronal coverage area

From the 30 µm brain sections, every 10^th^ section was collected for volumetric analysis. Sections were mounted on microscope slides and allowed to dry before staining for 10 minutes with 1% Sudan Black (S-2380, Sigma-Aldrich) made in 70% ethanol. Next, slides were washed three times with 70% ethanol and three times with PBS and then coverslipped using ProLong Gold Mounting Medium. The stained slices were imaged on a Keyence BZ-X Microscope at x10 magnification. In ImageJ, hippocampal area was measured for each cross-section by outlining the hippocampus. Overall volume was calculated using the following formula: volume = distance between sections (30µm per section x 10 sections in between stained sample sections = 0.3mm) x sum of areas of all stained sections. Hippocampal and posterior lateral volumes were measured across 7 sections, approximately between coordinates AP = -1.2mm and AP = -3.4mm with ImageJ.

For neuronal layer thickness, two brain sections were stained with primary antibody NeuN (ABN90, Milipore Sigma, 1:500 dilution) as described above. Briefly, sections were imaged at x40 magnification using FV300 confocal microscope. The thickness of DG granular cell layer of the hippocampus was measured on ImageJ software by taking percent area coverage.

### Primary cell culture

Primary microglia were prepared from neonatal mice at P2-3 from PS19/E4 mice as previously described^56^. Briefly, the cortices were separated from the meninges and minced with a sterile razor blade. The tissue was digested with 2.5% trypsin + DNase for 25 minutes at 37 °C. After incubation, the trypsin was neutralized with 30% FBS in DMEM and triturated until cells formed a uniformed suspension. The cells were spun at 300g for 10 min and the pellet was resuspended in 10% FBS in DMEM. For individual preparation, cells from each pup were plated onto PDL coated T25 flasks supplemented with 50ng/mL GM-CSF. Tail clippings were saved from each pup for genotyping. On day 4, fresh media was added onto the cells. On Day 10-12, the flasks containing mixed glia were placed onto an incubated shaker at 37 °C for 1 hour at 180rpm to shake off the microglia. After spinning the supernatant from the flasks for 10min at 300g, the resulting pellet was resuspended in complete media + 50ng/mL GM-CSF and plated on to poly-D-lysine (PDL) coated coverslips in a 24-well plate at 100k cells/well or added onto Day 4-5 neurons (see co-culture section). Oligodendrocyte precursor cells (OPCs) were isolated as previously described^40^. Briefly, the same flasks containing the mixed glial cells were placed back in the incubated shaker at 37 °C for 6 hour at 250rpm to shake off the OPCs. The collected media was then placed onto an untreated flask for 1 hour to allow microglia and astrocytes to settle, while the OPCs remained in suspension. The media was then collected and plated at 50k cells/well onto the Day 4-5 neurons or as a monoculture.

Primary neurons were prepared from neonatal mice at P0 from PS19/E4 or PS19/E4-Syn1 mice as previously described^57^. Briefly, the cortices were separated from the meninges and minced with a sterile razor blade. The tissue was digested with 5.5units/mL of papain for 13min at RT, then neutralized with trypsin inhibitor. The cells were washed with an optimum/glucose solution. The dissociated cells were plated on poly-L-lysine (PLL) coated coverslips 400k cells/mL in neurobasal + B27 100u/mL penicillin G, 100ug/mL streptomycin, and 1% GlutaMAX. The cells were fed with half media changes every 2-3 days, until Day 4-5 when the primary microglia were added.

### Primary co-culture of neurons, microglia, and OPCs

Primary neurons and microglia were co-cultured with both conditioned media as well as a physical co-culture. For the conditioned media culture, neuronal conditioned media from PS19/E4 and PS19/E4/Syn1-Cre neurons were collected from Day 7-10 neurons, then mixed 1:1 with microglia complete media and added onto the microglia. For the physical co-culture, isolated Day 10-12 microglia, and OPCs for tri-culture system, were plated on top of Day 7 primary neurons to form the co-culture for 72 hours. The media used for this co-culture was a 1:1 mixture of neurobasal B27 and microglia complete media supplemented with 50ng/mL of GM-CSF. After 72 hours, the cells were fixed with 4% PFA in DPBS for 15 minutes to be used for further analysis.

### BODIPY C-16 lipid transfer assay

For the lipid transfer assay, Day 10 live neurons were treated with media containing 1mg/mL BODIPY FL-C_16_ (4,4-Difluoro-5,7-Dimethyl-4-Bora-3a,4a-Diaza-*s*-Indacene-3-Hexadecanoic Acid) (D3821, Invitrogen, 1:1,000 dilution) for 1 hour at 37 °C. The cells were then washed with neurobasal B27 three times, before adding on the microglia for 24-hour co-culture treatment. After 24-hour co-culture, the cells were fixed as 4% PFA as described above then processed for ICC.

### In vitro immunocytochemistry (ICC) and BODIPY staining

Primary cells were plated onto 12-mm coated glass coverslips in 24-well plates. Coverslips were coated with PLL for neurons and PDL for microglia. Cells were fixed in 4% PFA for 15min, then washed 3x with 1x DPBS (14080055, Gibco). The rest of the ICC staining protocol is similar to the IHC described above with DPBS substituting for PBS. Briefly, cells were permeabilized during 1h block with 5% NDS in 0.2% Triton-X and one drop of M.O.M. for every 4mL of DPBS. After blocking, cells were stained with the following primary antibodies diluted to optimal concentrations overnight at 4C: microglia conditioned media group were stained for the following proteins: anti-PLIN2 (AB52356, Abcam, 1:1,000); anti-Iba1(rb) (AB178846, Abcam, 1:1,000); anti-Iba1 (gp) (234308, Synaptic Signals, 1:500); anti-Iba1 (gt) (AB5076, Abcam, 1:1,000); Anti-βIII Tubulin (ms) (G7121, Promega, 1:2,000); and Anti-βIII Tubulin (rb) (802011, BioLegend, 1:1,000); and anti-MBP (rt) (AB7349, Abcam, 1:200). Secondary antibodies were incubated in: donkey anti-rabbit 405 (AB175651, Abcam, 1:1,000); donkey anti-rabbit 488 (A21206, Invitrogen, 1:1,000); donkey anti-mouse 594 (A21203, Invitrogen, 1:1,000); donkey anti-rabbit 594 (A21207, Invitrogen, 1:1,000); donkey anti-rat 594 (A21209, Invitrogen, 1:1,000); donkey anti-rabbit 647 (A31573, Invitrogen, 1:1,000); and donkey anti-guinea pig 647 (706-65-148, Jackon Immuno, 1:1,000 dilution). Coverslips were mounted to microscope slides with VECTASHIELD Prolong Gold with DAPI (H-1200-10, Vectors Labs) or without DAPI if the 405 channel was occupies (P36930, Vector Labs). Images were taken with FV3000 confocal laser scanning microscope (Olympus) at ×20, ×40, or 860 magnification depending on the stain. BODIPY and BODIPY C11 staining were performed as described in the IHC section.

### Quantitative analysis of immunohistological and immunocytochemical data

For quantifications, two sections (∼300 µm apart) from each brain were selected from each animal and processed for immunohistochemistry. DG endpoints were analyzed with ImageJ software. Percent area coverage of various cell types was analyzed by manually drawing a ROI (region of interest) around the DG hilus, including the neuronal layer. To quantify percent area coverage of lipids marked by PLIN2^+^ or BODIPY^+^ inside specific cell types, a mask was first created with a marker from that cell type. The mask was then applied to the lipid stain and subsequent percent area coverage was measured. For example, to analyze the percent area of BODIPY^+^ neutral lipids inside Iba1^+^ microglia, a mask was created from the Iba1^+^ channel, then applied to the BODIPY^+^ channel. To assess the percent area coverage of BODIPY in the DG, the hilus area of the DG, including the neuronal layer, was manually dawned identified via DAPI staining in 40x images and the BODIPY percent area of that ROI was calculated.

For the percent coverage area quantification, an optimal threshold was established for each stain in ImageJ and all samples were quantified utilizing the established threshold and automated ImageJ macros for each stain. For percent count when analyzing the percentage of Olig2^+^ and BODIPY^+^ markers inside Iba1^+^ microglia, images were counted with cell count tool on ImageJ. To exclude the possibility of bias, researchers were blinded to samples during analysis.

### Single-nuclei preparation for 10x loading

The snap-frozen dissected mouse hippocampus were placed on ice and placed into a pre-chilled 2 ml Dounce with 1 ml of cold 1× Homogenization Buffer (1× HB) (250 mM sucrose, 25 mM KCl, 5 mM MgCl_2_, 20 mM tricine-KOH pH 7.8, 1 mM dithiothreitol, 0.5 mM spermidine, 0.15 mM spermine, 0.3% NP40, 0.2 U µl^−1^ RNase inhibitor and ∼0.07 tablets per sample protease inhibitor). Dounce with ‘A’ loose pestle (strokes) and then with ‘B’ tight pestle (10 strokes). The homogenate was filtered using a 70-µm Flowmi strainer (VWR) and transferred to a pre-chilled 2-ml LoBind tube (Fisher Scientific). Nuclei were pelleted by spinning for 5 min at 4 °C at 350 RCF. The supernatant was removed and the nuclei were resuspended in 400 µL 1X HB in a pre-chilled LoBind tube. Next, 400 µl of 50% Iodixanol solution was added to the nuclei and then slowly layered with 600 µl of 30% Iodixanol solution under the 25% mixture, then layered with 600 µl of 40% Iodixanol solution under the 30% mixture. The nuclei were then spun for 20 min at 4°C at 3,000 RCF in a pre-chilled swinging bucket centrifuge. Then 200 µl of the nuclei band at the 30%–40% interface was collected and transferred to a fresh tube. Then, 800 µl of 2.5% BSA in PBS plus 0.2 U µl^−1^ of RNase inhibitor was added to the nuclei and then were spun for 10 min at 500 RCF at 4 °C. The nuclei were resuspended with 2.5% BSA in PBS plus 0.2 U µl^−1^ RNase inhibitor to reach ∼1,000-2,000 nuclei per µl. The nuclei were then filtered with a 40-µm Flowmi strainer. The nuclei were counted and then ∼14,000 nuclei per sample were loaded onto 10x Genomics Next GEM chip G. The snRNA-seq libraries were prepared using the Chromium Next GEM Single Cell 3ʹ Library and Gel Bead kit v.3.1 (10x Genomics) according to the manufacturer’s instructions. Libraries were sequenced on an Illumina NovaSeq X Plus sequencer at the UCSF CAT Core.

### Pre-processing and Clustering of Mouse snRNA-seq Samples

The snRNA-seq dataset included 16 total samples, comprising four mice per genotype group (PS19/E3, PS19/E4, PS19/NSE-E4, PS19/E4/Syn1-Cre). Each group consisted of 2 male and 2 female mice. The demultiplexed FASTQ files were aligned to a custom mouse reference genome using the 10x Genomics Cell Ranger Count pipeline^58^ (version 9.0.1), following the Cell Ranger documentation. The include-introns flag was set to TRUE to capture reads mapping to intronic regions.

The filtered count matrices generated by the Cell Ranger count pipeline for the 16 samples were processed using the R package Seurat^59^ (version 4.4.0) for single-nucleus analysis. Each sample was pre-processed as a Seurat object, then all 16 samples were merged into a single Seurat object. Low-quality nuclei were removed using the following criteria: fewer than 500 total UMI counts, fewer than 200 features, or a mitochondrial gene ratio higher than 0.25%. Sparse gene features expressed in fewer than 10 cells were also excluded. After quality control, normalization and variance stabilization was performed using SCTransform^60^ (version 0.4.1) method for initial parameter estimation. Graph-based clustering was conducted using the Seurat functions, FindNeighbors and FindClusters. Cells were embedded in a k-nearest neighbor (KNN) graph on the first 50 principal components (PCs) using Euclidean distance in the PCA space, and edge weights between two cells were refined using Jaccard similarity. Clustering via the Louvain algorithm (FindClusters) with 15 PCs and a resolution of 0.7 produced 36 distinct biologically relevant clusters, which were subsequently used for further analyses.

### Cell type assignment

Cell type identities were annotated following a method we previously described^61,62^. Briefly, Seurat’s AddModuleScore() function was applied to the object using features from lists of known brain marker genes (up to 8 per cell type), which were previously reported in a study^57^ that provided a publicly available resource of brain-wide in situ hybridization images. Further subdivisions of hippocampus cell types, such as CA1 and CA3 pyramidal cells, were identified using hippocampus-specific marker genes curated from Hipposeq (https://hipposeq.janelia.org). A module score was computed for each cell type in every cell, and each cell was assigned the cell type that produced the highest module score among all tested cell types. Cells with their top two scores within 10% of each other were classified as potential hybrids and excluded from downstream analyses. Cells with negative scores for all cell types computed were classified as negative and excluded from downstream analyses. The cell type assignments per cluster was evaluated by inspecting homogeneity, distribution, and spatial separation of cluster in UMAP space. Each cluster was annotated based on its dominated cell type identity. Clusters containing comparable proportions of two or more cell types were deemed unresolved. Dot-plot of canonical marker genes were generated using the DotPlot() function in Seurat (version 4.4.0).

### Gene set enrichment analysis

DE genes between clusters of interest or between two genotype groups were identified using the FindMarkers Seurat function on the SCT assay data. This algorithm uses the Wilcoxon rank-sum test to identify DE genes between two populations. DE genes were limited to genes detected in at least 10% of the cells in either population and with at least 0.1 log_2_ fold change. Volcano plots with log_2_ fold change and *P* value from the DE gene lists were generated using the ggplot2 R package (version 3.5.2). Overrepresentation (or enrichment) analysis was performed using clusterProfiler^63^ (version 4.14.6) to find gene sets associated with the DE genes with at least 10 genes in the mouse-specific KEGG database^64^. The *P* values are based on a hypergeometric test and are adjusted for multiple testing using the Benjamini–Hochberg method^65^. The same method was used for gene set enrichment analysis of astrocyte subclusters and microglia subclusters.

### Association between clusters and genotype

A GLMM_AM was implemented in the lme4 (version 1.1-37) R package^66^ and used to estimate the associations between cluster membership and the mouse model. These models were run separately for each cluster of cells. The GLM model was performed with the family argument set to the binomial probability distribution and with the ‘nAGQ’ parameter set to 10 corresponding to the number of points per axis for evaluating the adaptive Gauss–Hermite approximation for the log-likelihood estimation. Cluster membership of cells by sample was modeled as a response variable by a two-dimensional vector representing the number of cells from the given sample belonging to and not belonging to the cluster under consideration. The corresponding mouse ID from which the cell was derived was the random effect variable, and the animal model for this mouse ID was included as the fixed variable. The reference animal model was set to PS19/E4. The resulting *P* values for the estimated LOR across the four animal models (with respect to the PS19/E4) and clusters were adjusted for multiple testing using the Benjamini–Hochberg method^65^. The same method was used for estimating the between-cluster association with genotype for astrocyte subclusters and microglia subclusters.

### Association between proportion of cell types and histopathological parameters

A GLMM_histopathology (GLMM_LP for this study) was implemented in the lme4 (version 1.1-37) R package^66^ and used to identify cell types whose proportions are significantly associated with changes in histopathology across the samples. These models were performed separately for each combination of all clusters and one histological parameters. The GLM model was performed with the family argument set to the binomial probability distribution family and with the ‘nAGQ’ parameter set to 1 corresponding to a Laplace approximation for the log-likelihood estimation. Cluster membership of cells by sample was modeled as a response variable by a two-dimensional vector representing the number of cells from the given sample belonging to and not belonging to the cluster under consideration. The corresponding mouse model from which the cell was derived was included as a random effect, and, further, the mouse ID within the given mouse model was modeled as a random effect as well. Note, this represents the hierarchical nature of these data for the GLMM, and the mouse models are first assumed to be sampled from a ‘universe’ of mouse models; this is then followed by sampling mice within each mouse model. The modeling choice of including the mouse model as a random effect as opposed to a fixed effect is meant to increase the degrees of freedom (or maximize the statistical power) to detect the association of interest, particularly in light of the relatively small number of replicates (3–4) per animal model. The histological parameter under consideration was modeled as a fixed effect in this model.

We selected a subset of cell types of interest and visualized the LOR estimates (derived from the GLMM fits) in a heat map using pheatmap package 1.0.12 after adjusting the *P* values distribution across histopathological parameters across cell types with Benjamini–Hochberg multiple testing correction^65^.

We applied the pipeline to the astrocyte and microglia subtypes and visualized the associations between astrocyte and microglia subtypes of interest and the histopathological parameters.

### General Statistics and Reproducibility

Sample sizes for mouse studies were chosen on the basis of estimates to provide statistical power of ≥80% and alpha of 0.05 based on preliminary data. For analysis with multiple groups, the differences between genotype groups were evaluated by ordinary one-way ANOVA with Tukey’s multiple comparisons test, where the mean of each column was compared to the mean of every other column. For analysis with two groups, the differences between the genotypes were evaluated by unpaired two-sided t-test. All data are shown as mean ± s.e.m. Data distribution was assumed to be normal, though this was not formally tested. For correlations between two data sets, simple linear regression or nonlinear regression R^2^ were used. p < 0.05 was considered to be significant, and all significant p values were included in figures or noted in figure legends. Statistical significance analysis and plots were completed with GraphPad Prism 10 for Mac (GraphPad Software). No randomization method was used for the assignment of mice to study groups, and no animals or data points were excluded from these studies.

For mouse snRNA-seq studies, sample sizes were determined by a power analysis based on effect sizes from our previous studies and literature. Mice selected for snRNA-seq consisted of 2 males and 2 females per group that together represented average pathologies for all parameters quantified. Nuclei were isolated from four mice per genotype to ensure n ≥ 3 mice per group. Pathology data were correlated with snRNA-seq data. Investigators were not blinded during analysis of the snRNA-seq datasets, as sample metadata were needed to conduct any comparisons. Studies were all performed using one cohort of mice.

## Data availability

The H. sapiens MAPT sequence is available at https://www.ncbi.nlm.nih.gov/nuccore/NM_001123066. The reference mouse genome sequence (GRCm38) from Ensembl (release 98) is available at http://ftp.ensembl.org/pub/release-98/fasta/mus_musculus/dna/Musmusculus.GRCm38.dna.primary_assembly.fa.gz. The reference mouse gene annotation file from GENCODE (release M23) is available at http://ftp.ebi.ac.uk/pub/databases/gencode/Gencode_mouse/release_M23/gencode.vM23.primary_asse mbly.annotation.gtf.gz. The snRNA-seq datasets generated during the study are available at the Gene Expression Omnibus under accession number <INTENDED link>.

Data associated with Figure 5 and 6 and Extended Data Figure 7 and 8 are also available in the Supplementary Information. Source data are provided with this paper.

## Code availability

The following packages/software were used either as dependencies to downloading or using packages mentioned in the Methods section or in creating the figures in this manuscript: Seurat v.4.4.0, CellRanger v.9.0.1, glm lme4 v.1.1-37, clusterProfiler v.4.14.6, ggplot2 v.3.5.2, pheatmap v.1.0.12, and Fiji v.2.1.0 (ImageJ). GraphPad Prism v.10.6.1 was used for some statistical analyses and graphing. All codes generated with custom R and shell scripting during this study are accessible via GitHub at <INTENDED github link for paper>.

## Acknowledgements

This work was partially supported by National Institutes of Health (NIH) grants R01AG071697, RF1AG076647, and P01AG073082 to Y.Huang. The funders had no role in study design, data collection and analysis, decision to publish or preparation of the manuscript. We thank the entire Huang laboratory and Gladstone Institutes of Neurological Disease (GIND) colleagues for critical discussion and feedback; R. Bladino of the Gladstone Histology and Light Microscopy Core for advice; the UCSF Cat core, for sequencing, supported by UCSF PBBR, RRP IMIA, and NIH 1S10OD028511-01 grants; and T.Pak for editorial assistance.

## Author Contributions

M.J.K. and Y.Huang collaborated on the overall study design, coordinated the experiments, and wrote the manuscript. M.J.K. performed most of the studies, including all primary culture studies, immunofluorescent stains, data collections and analysis, and generated all the figures. J.B., D.S., and N.K. helped with mouse sample collection. J.B. performed analysis for hippocampal volume and p-tau for all PS19/NSE-E4 and some of the PS19/E4, PS19/E3, PS19/E4/Syn1-Cre mice. O.Y. and S.A. helped with some of the primary culture preparations. Z.P. and Z.L helped with some of the immunohistochemistry, p-tau, and hippocampal volume analysis. Z.P. and S.A. helped with some of the confocal imaging. Y.Hao and K.S. isolated cell nuclei and prepared samples for snRNA-seq. Y.L. and L.Y. performed the snRNA analysis. S.D.L. set up breeding cages and performed genotyping for primary culture studies. J.N. helped with setting up breeding cages and feeding PLX5622 study mice. C.E. managed all mouse lines. Y.Huang supervised the project.

---

**Extended Data Fig. 1.**
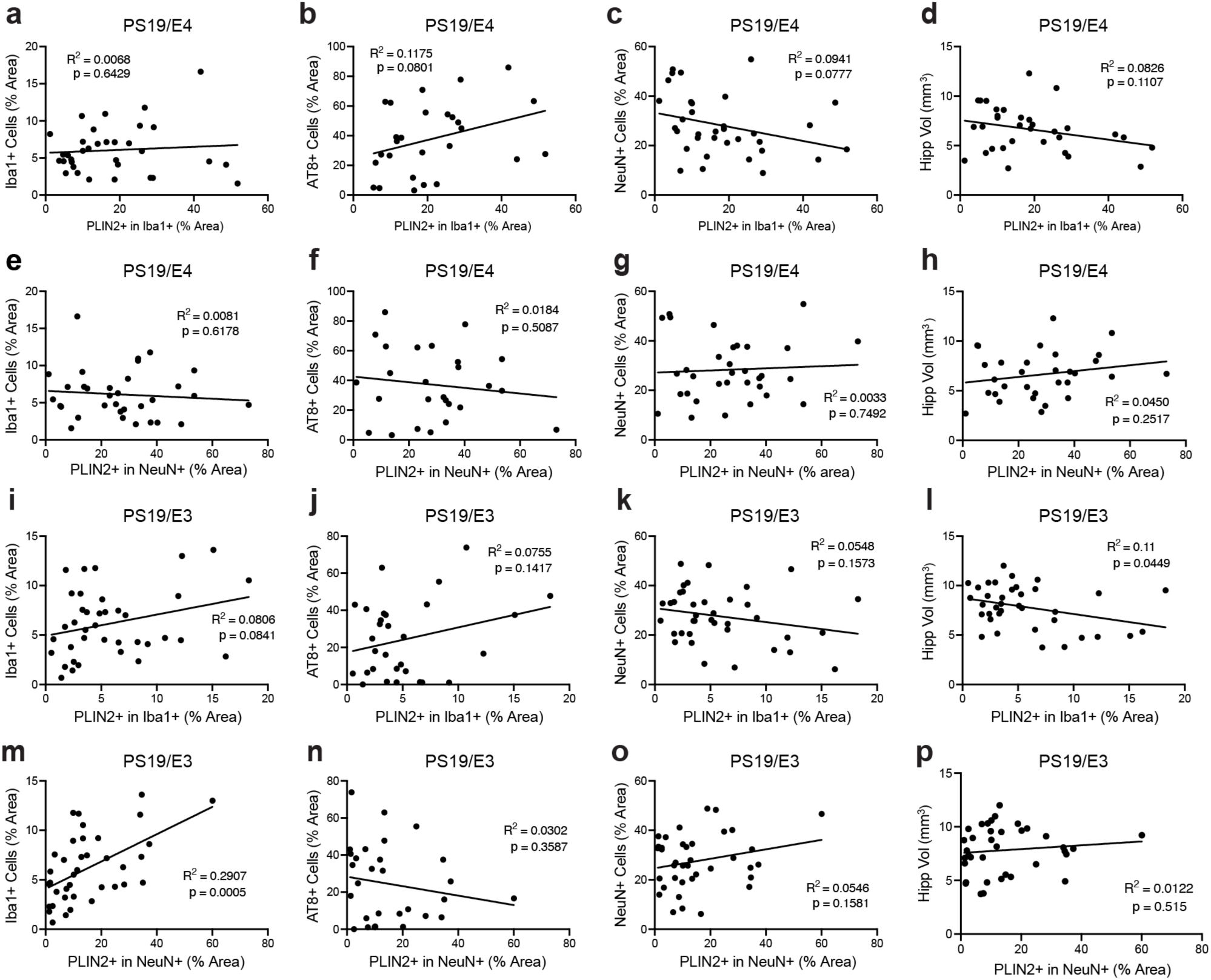
| Lipid droplet accumulation shows only mild correlation with tau induced pathologies in PS19/E4 and PS19/E3 mice. a-d, Correlations between PLIN2^+^ lipid droplet (LD) % area in Iba1^+^ microglia and Iba1^+^ % area in the hippocampus (*n*= 34) (**a**), AT8^+^ % area in the hippocampus (*n*= 34) (**b**), NeuN^+^ % area in the hippocampus (*n*= 35) (**c**), and hippocampal volume (*n*= 33) (**d**) in PS19/E4 mice (*n*= 35). **e-h,** Correlations between PLIN2^+^ LD % area in NeuN^+^ neurons and Iba1^+^ % area (**e**), AT8^+^ % area (**f**), NeuN^+^ % area (**g**), and hippocampal volume (**h**) in PS19/E4 mice. **i-l,** Correlations between PLIN2^+^ LD % area in Iba1^+^ microglia and Iba1^+^ % area (*n*= 39) (**i**), AT8^+^ % area (*n*= 31) (**j**), NeuN^+^ % area (*n*= 39) (**k**), and hippocampal volume (*n*= 39) (**l**) in PS19/E3 mice. **m-p,** Correlations between PLIN2^+^ LD % area in NeuN^+^ neurons and Iba1^+^ % area (**m**), AT8^+^ % area (**n**), NeuN^+^ % area (**o**), and hippocampal volume (**p**) in PS19/E3 mice (*n*= 42). Pearson’s correlation analysis (two-sided) was used in **a-p**.

**Extended Data Fig. 2.**
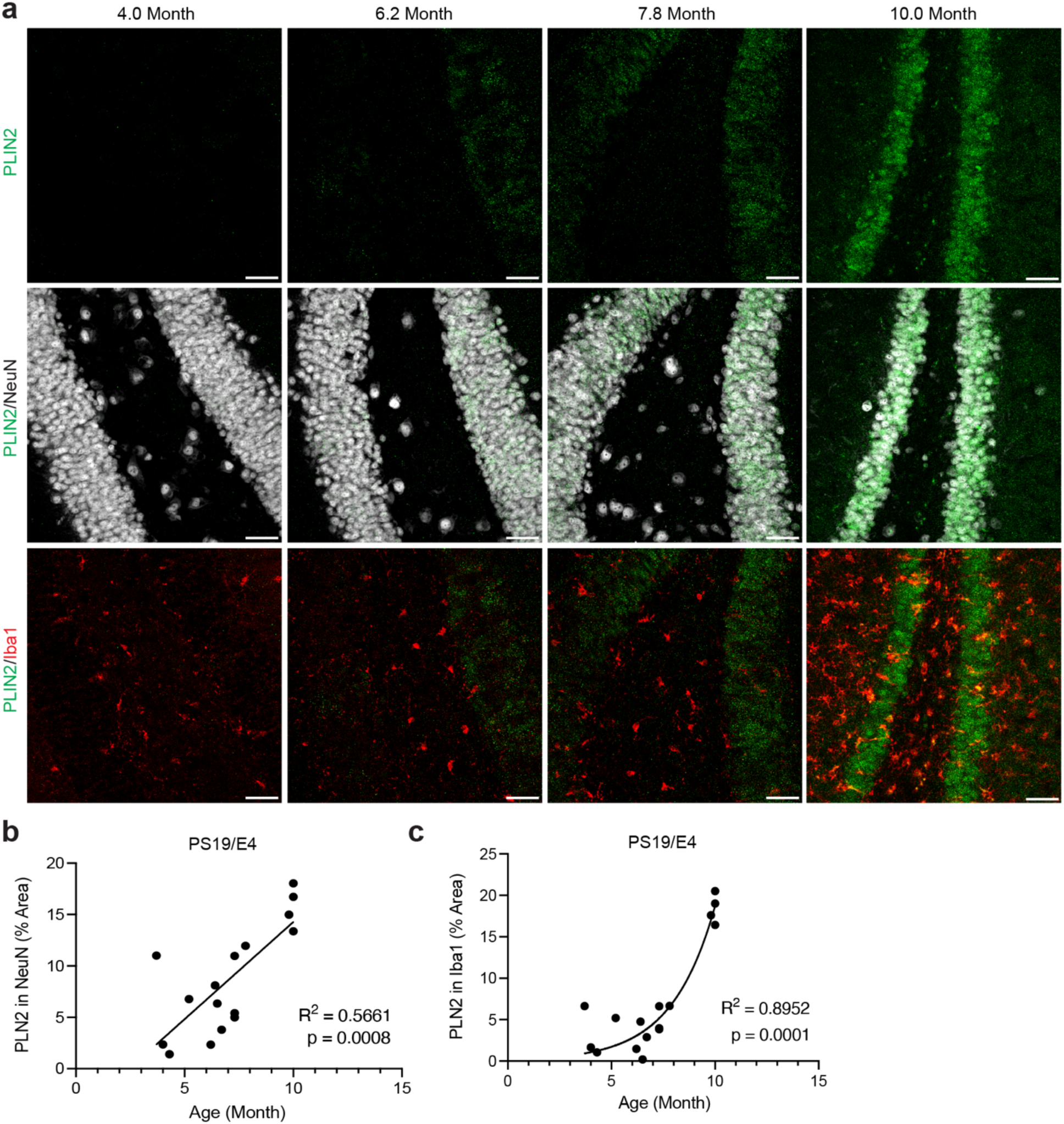
| Neuronal lipid droplets (LDs) accumulate linearly with age while microglial LDs accumulate exponentially at the gliosis onset in PS19/E4 mice. **a,** Representative immunofluorescent images of PLIN2^+^ LD (green), NeuN^+^ neurons (white), and Iba1^+^ microglia (red) in PS19/E4-mice at 4.0, 6.2, 7.8, and 10.0 months of age. Scale bars, 40μm. **b,** Correlation between age (in months) and PLIN2^+^ LDs in NeuN^+^ neurons. Pearson’s correlation analysis (two-sided). **c,** Correlation between age (in months) and PLIN2^+^ LDs in Iba1^+^ microglia. Nonlinear regression R^2^. All scale bars, 40μm. For **b,c** PS19/E4 mice, *n* = 16

**Extended Data Fig. 3.**
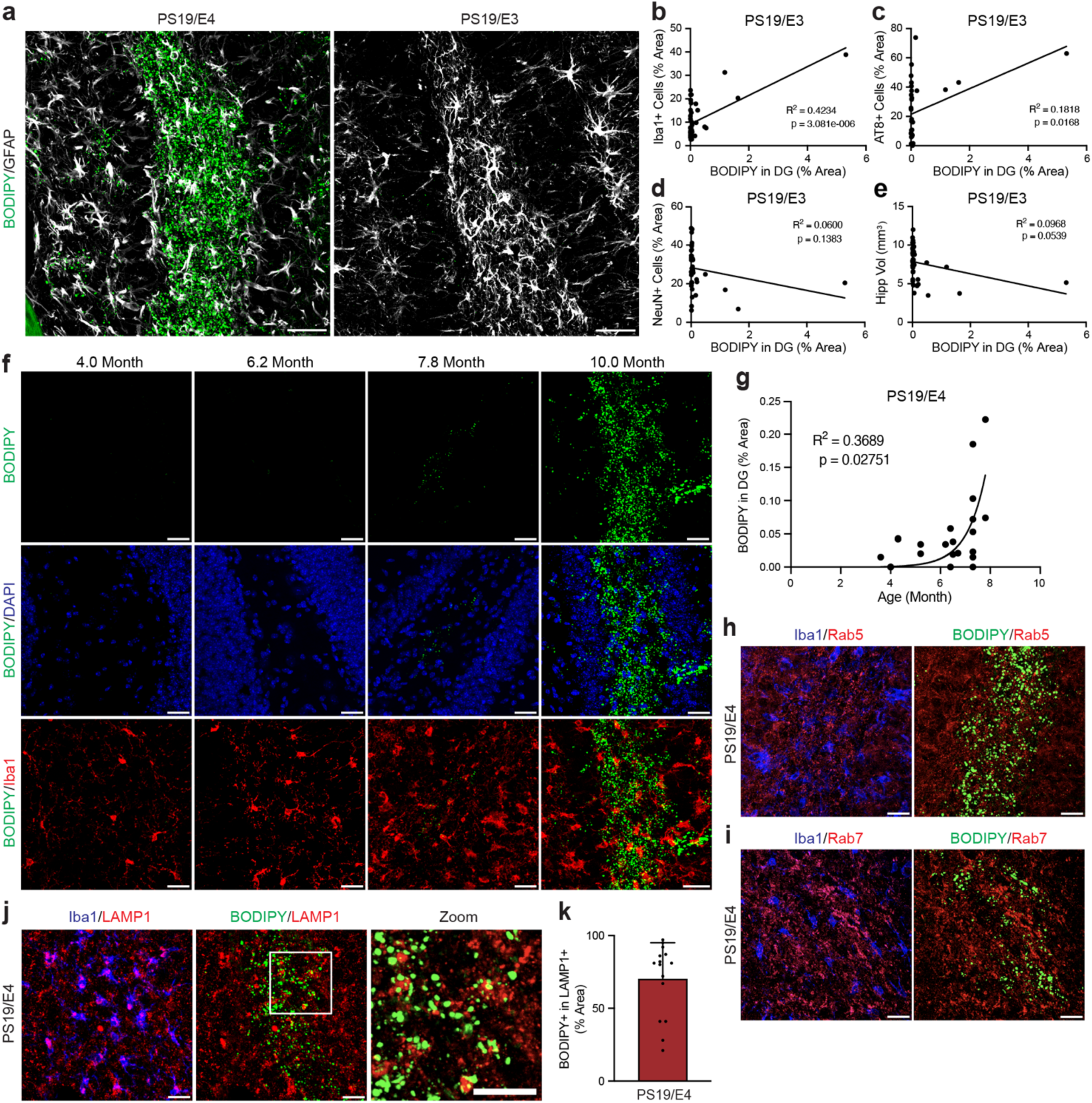
| Neutral lipids accumulation in the hippocampal dentate gyrus (DG) of PS19/E4 mice and their subcellular localization in microglia. **a,** Representative immunofluorescent images of BODIPY^+^ neutral lipids and GFAP^+^ astrocyte in the hippocampal DG of PS19/E4 and PS19/E3 mice at 10 months of age. **b-e,** Correlations between BODIPY^+^ neutral lipids in DG (% area) and Iba1^+^ % area (*n*= 42) (**b**), AT8^+^ % area (*n*= 31) (**c**), NeuN^+^ % area (*n*= 39) (**d**), and hippocampal volume (*n*= 39) (**e**) in PS19/E3 mice, assessed via Pearson’s correlation analysis (two-sided). **f,** Representative immunofluorescent images of BODIPY^+^ neutral lipids (green), DAPI^+^ nuclei (white), and Iba1^+^ microglia (red) in the hippocampal DG of PS19/E4 mice at 4.0, 6.2, 7.8, and 10.0 months of age. Scale bars, 25μm. **g,** Correlation between age (in months) and BODIPY^+^ neutral lipids in the hippocampal DG of PS19/E4 mice (*n*= 20). Nonlinear regression R^2^. **h-j,** Representative immunofluorescent images of Rab5^+^ early endosomes (red, **h**), Rab7^+^ late endosomes (red, **i**), and LAMP1^+^ lysosomes (red, **j**) with Iba1^+^ microglia (blue) and BODIPY^+^ neutral lipid (green) in PS19/E4 mice. All scale bars, 40μm. **k,** Quantification of BODIPY^+^ marker inside LAMP1^+^ stain (*n*= 15).

**Extended Data Fig. 4.**
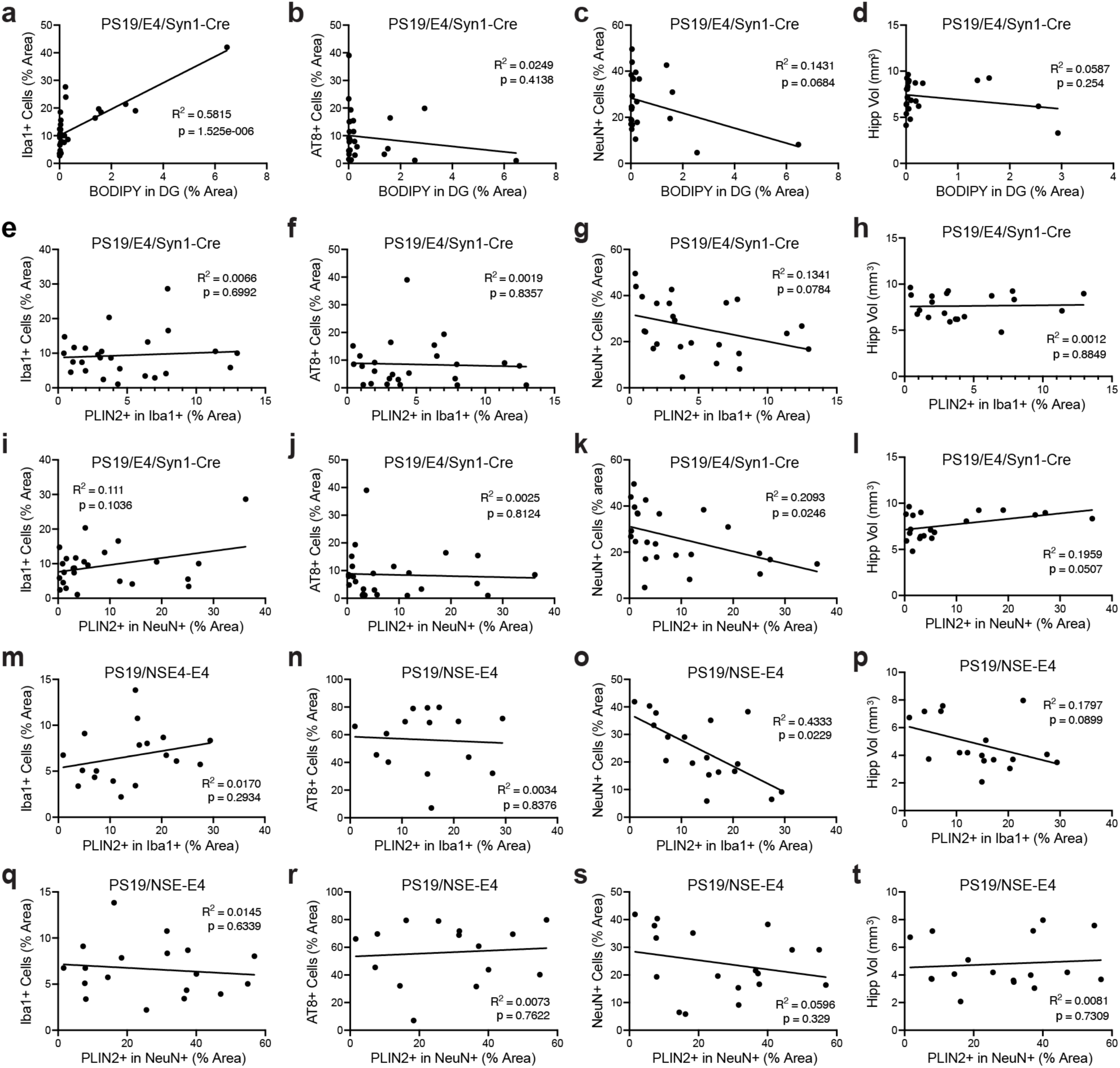
| Lipid droplet accumulation shows only mild correlation with tau-induced pathologies in PS19/E4-Syn1-Cre and PS19/NSE-E4 mice. a-d, Correlations between BODIPY^+^ neutral lipids in DG (% area) and Iba1^+^ % area (*n*= 29) (**a**), AT8^+^ % area (*n*= 25) (**b**), NeuN^+^ % area (*n*= 29) (**c**), and hippocampal volume (*n*= 24) (**d**) in PS19/E4/Syn1-Cre mice. **e-h,** Correlations between PLIN2^+^ lipid droplets (LDs) (% area) in Iba1^+^ microglia and Iba1^+^ % area (*n*= 25) (**e**), AT8^+^ % area (*n*= 25) (**f**), NeuN^+^ % area (*n*= 24) (**g**), and hippocampal volume (*n*= 24) (**h**) in PS19/E4/Syn1-Cre mice. **i-l,** Correlations between PLIN2^+^ LDs (% area) in NeuN^+^ neurons and Iba1^+^ % area (*n*= 25) (**i**), AT8^+^ % area (*n*= 25) (**j**), NeuN^+^ % area (*n*= 24) (**k**), and hippocampal volume (*n*= 24) (**l**) in PS19/E4/Syn1-Cre mice. **m-p,** Correlations between PLIN2^+^ LDs (% area) in Iba1^+^ microglia and Iba1^+^ % area (*n*= 29) (**m**), AT8^+^ % area (*n*= 15) (**n**), NeuN^+^ % area (*n*= 18) (**o**), and hippocampal volume (*n*= 17) (**p**) in PS19/NSE-E4 mice (*n*= 18). **q-t,** Correlations between PLIN2^+^ LDs (% area) in NeuN^+^ neurons and Iba1^+^ % area (*n*= 18) (**q**), AT8^+^ % area (*n*= 15) (**r**), NeuN^+^ % area (*n*= 18) (**s**), and hippocampal volume (*n*= 17) (**t**) in PS19/NSE-E4 mice. Pearson’s correlation analysis (two-sided) was used for **a-t**.

**Extended Data Fig. 5.**
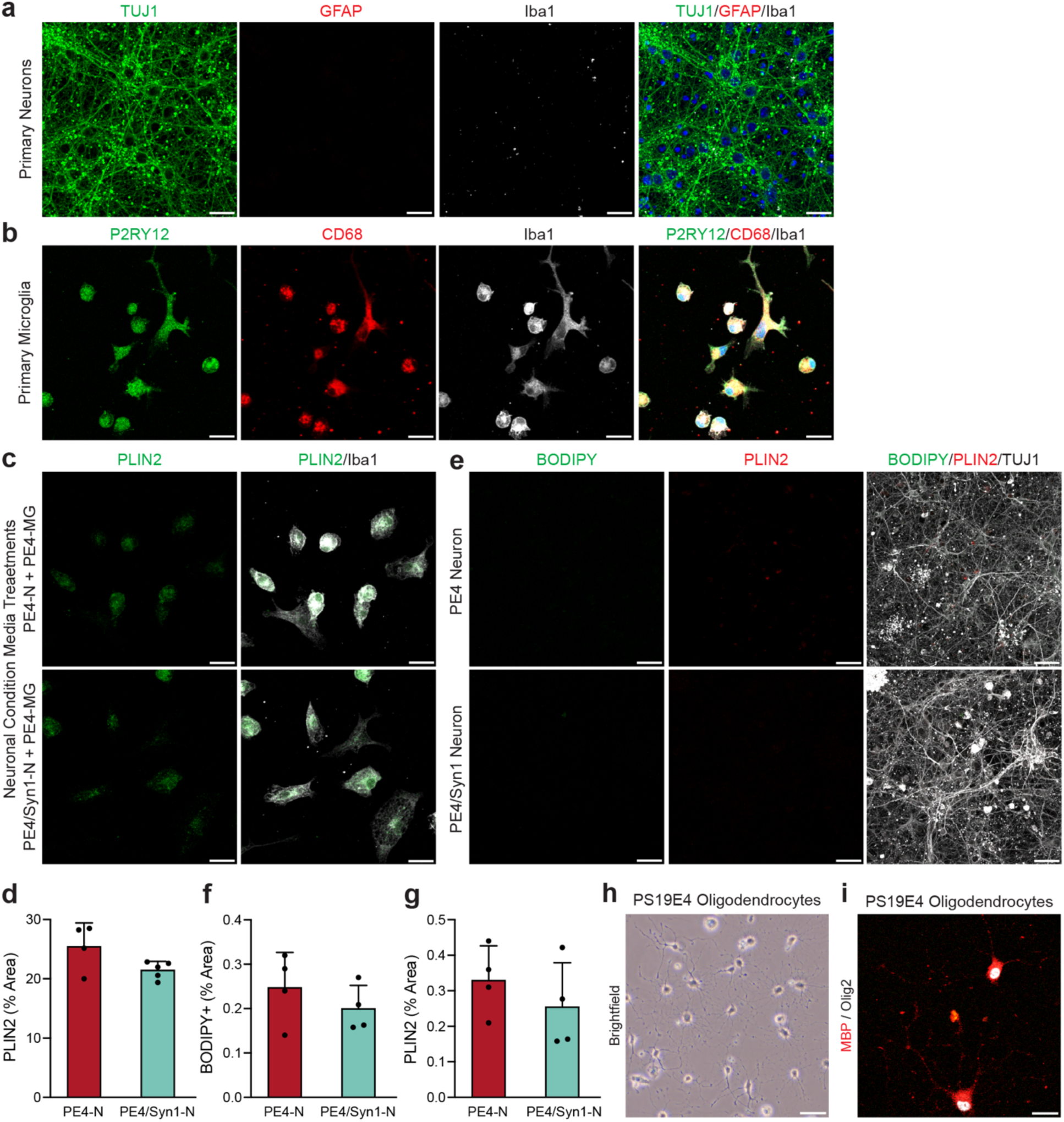
| Validation of primary neuron and glia co-cultures. **a,** Representative immunofluorescent images of TUJ1^+^ neurons (green), GFAP^+^ astrocytes (red), and Iba1^+^ microglia (white) in primary mouse neuron culture. Iba1^+^ (white channel) shows minimal signal due to background debris. **b,** Representative immunofluorescent images of P2RY12^+^ microglia (green), CD68^+^ microglia (red), and Iba1^+^ microglia (white) in primary mouse microglia culture. **c,** Representative immunofluorescent images of PLIN2^+^ lipid droplets (LDs, green) in primary PS19/E4 Iba1^+^ microglia (white) treated with neuron-conditioned media (either PS19/E4 neuron CM or PS19/E4/Syn1-Cre neuron CM) for 72 hours. **d,** Quantification of percent PLIN2^+^ area coverage in Iba1^+^ microglia (PS19/E4 neuron CM, *n*= 4; PS19/E4/Syn1-Cre neuron CM, *n*= 4). **e,** Representative immunofluorescent images of BODIPY^+^ neutral lipids (green), PLIN2^+^ LDs (red) in TUJ1^+^ neurons (white) from PS19/E4 or PS19/E4/Syn1-Cre mice. **f,** Quantification of percent BODIPY^+^ area coverage in TUJ1^+^ neurons (PS19/E4 neuron, *n*= 4; PS19/E4/Syn1-Cre neuron, *n*= 5). **g,** Quantification of percent PLIN2^+^ area coverage in TUJ1^+^ neurons (PS19/E4 neuron, *n*= 4; PS19/E4/Syn1-Cre neuron, *n*= 5). **h,** Representative brightfield image of morphological observation of mature oligodendrocytes. Scale bar, 40μm. **i,** Representative immunofluorescent images of MBP^+^ (red) and Olig2^+^ (white) oligodendrocytes in PS19/E4. All scale bars, 10μm unless otherwise stated. Quantified data are represented in **d,f,g** as mean ± s.e.m., unpaired two-sided t-test.

**Extended Data Fig. 6.**
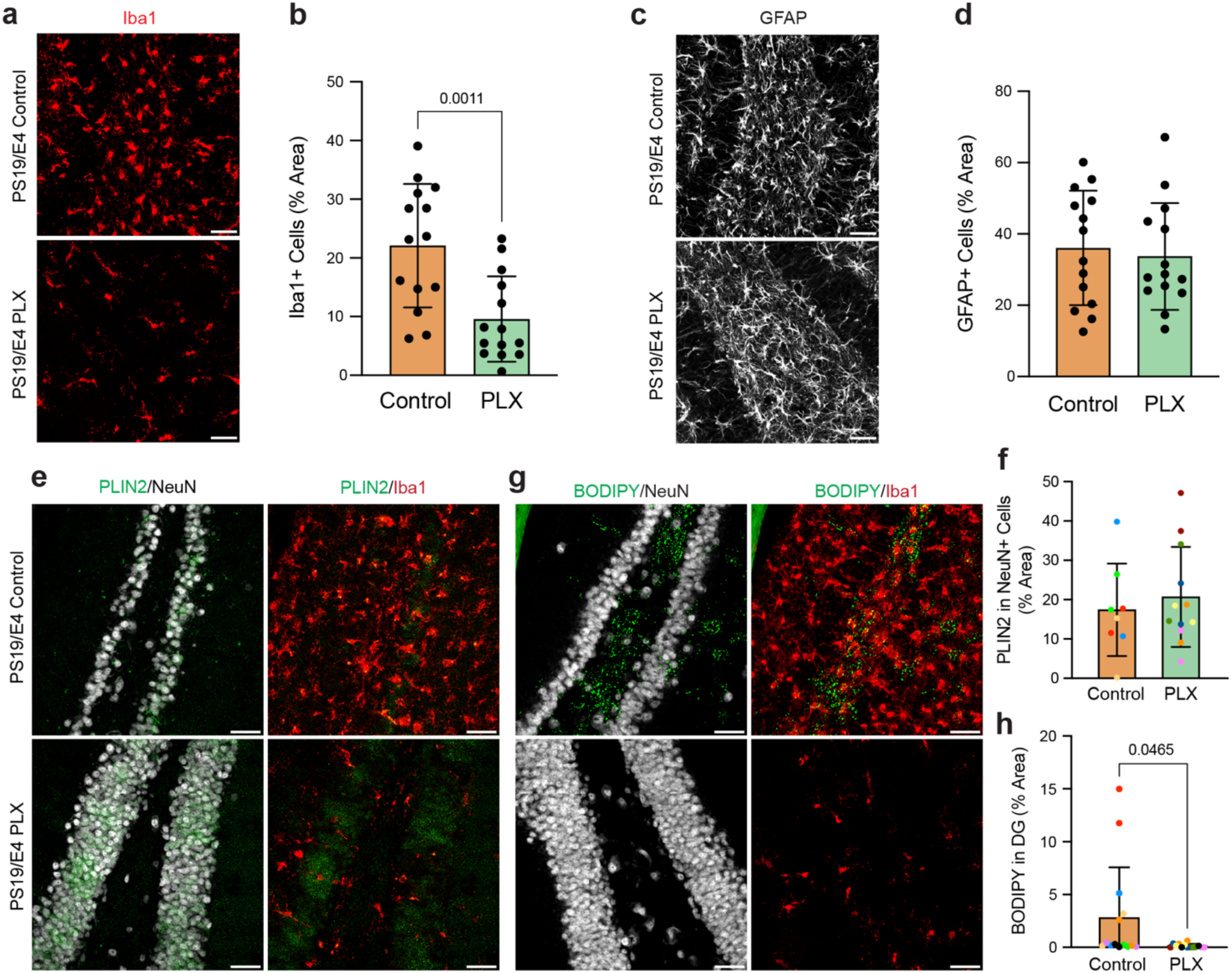
| Microglia are necessary for neutral lipid accumulation in the hippocampal dentate gyrus (DG) in PS19 tauopathy mice. **a,** Representative immunofluorescent images of Iba1^+^ microglia (red) in 10-month-old PS19/E4 mice treated with either control or PLX5622 (PLX) chow for 3 months. **b,** Quantification of percent Iba1^+^ area coverage in PS19/E4-Control or PS19/E4-PLX mice. **c,** Representative immunofluorescent images of GFAP^+^ astrocytes (white) in 10-month-old PS19/E4 mice treated with either control or PLX5622 chow for 3 months. **d,** Quantification of percent GFAP^+^ area coverage in PS19/E4-Control or PS19/E4-PLX mice. **e,** Representative immunofluorescent images of PLIN2^+^ lipid droplets (green), NeuN^+^ neurons (white), and Iba1^+^ microglia (red) in the hippocampal DG of 10-month-old PS19/E4-Control or PS19/E4-PLX mice. **f,** Quantification of percent PLIN2^+^ area coverage in Iba1^+^ microglia. **g,** Representative immunofluorescent images of PLIN2^+^ lipid droplets (green), NeuN^+^ neurons (white), and Iba1^+^ microglia (red) in the hippocampal DG of 10-month-old PS19/E4-Control or PS19/E4-PLX mice. **h,** Quantification of percent BODIPY^+^ area coverage in the hippocampal DG of PS19/E4-Control or PS19/E4-PLX mice. In **a,c,e,g,** all scale bars, 40μm. In **f,h,** each color corresponds to one mouse, with two brain sections per mouse for PS19/E4-Control mice (*n*= 4) and PS19/E4-PLX mice (*n=* 6). Quantified data in **b,d,f,h** are represented as mean ± s.e.m., unpaired two-sided t-test.

**Extended Data Fig. 7.**
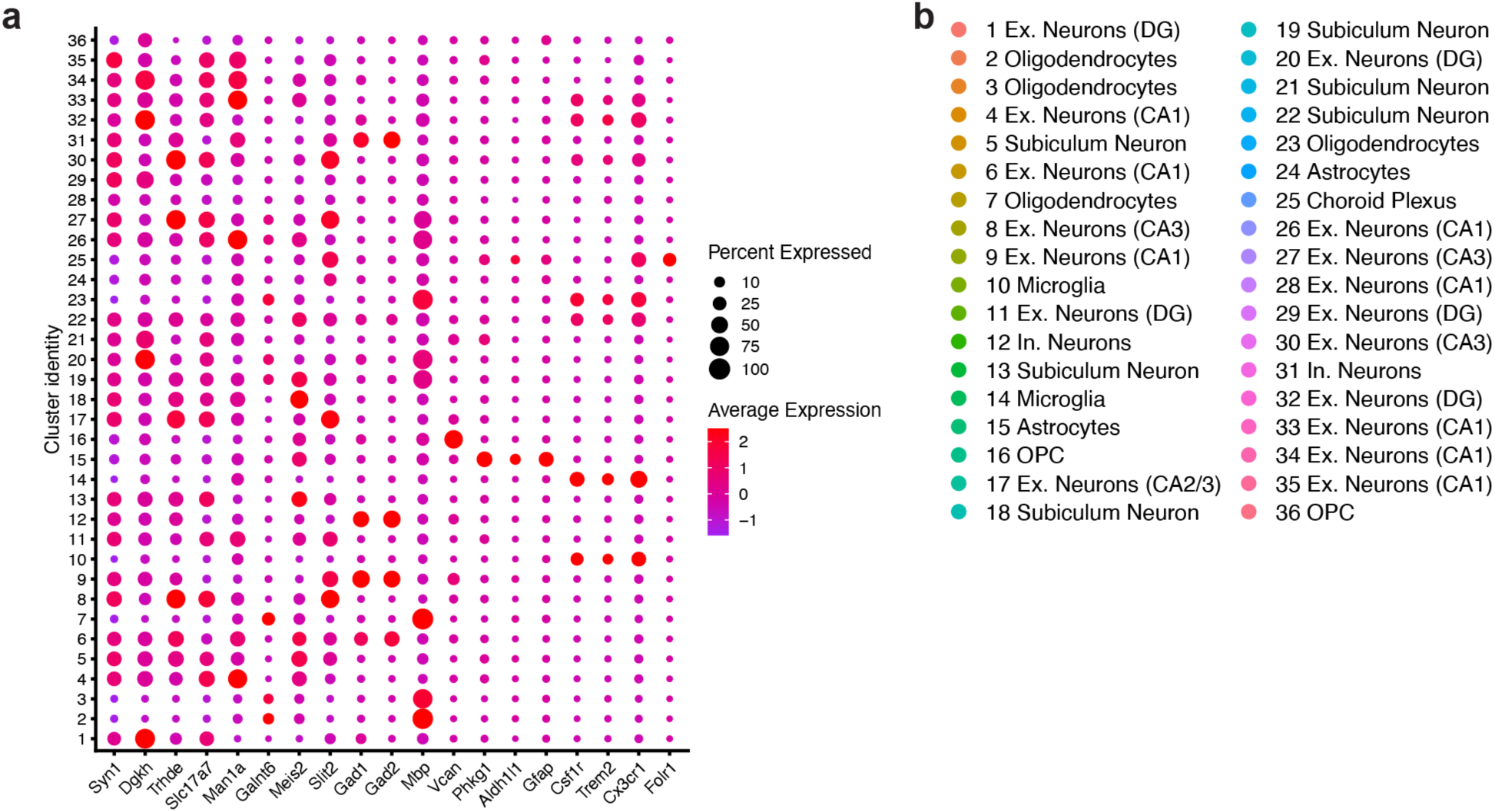
| Cell cluster identification by snRNA-seq analysis of tauopathy mice with different APOE genotypes. **a**, Dot-plot depicting normalized average expression of selected cell identity marker genes for all 36 distinct hippocampal cell clusters from mice with different APOE genotypes at 10 months of age. **b**, Cluster identity of 36 distinct cell types. Ex neuron, excitatory neuron; In neuron, inhibitory neuron.

**Extended Data Fig. 8.**
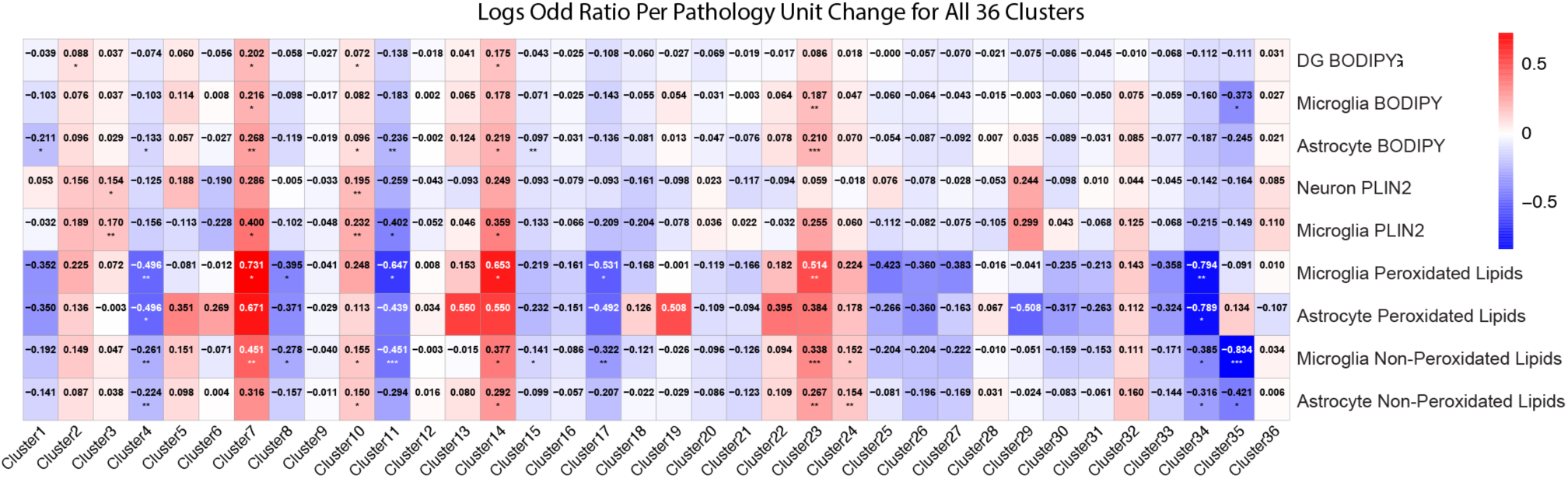
| Heat map plot of the log odds ratio per unit change in each pathological lipid parameter for all cell clusters. The LOR represents the mean estimate of the change in the log odds of cells per sample from a given animal model, corresponding to a unit change in a given histopathological parameter. Associations with pathologies are colored (negative associations, blue; positive associations, red). P values are from fits to a GLMM_histopathology; the associated tests are two-sided. * p < 0.05, ** p <0.01, *** p <0.001.

**Extended Data Fig. 9.**
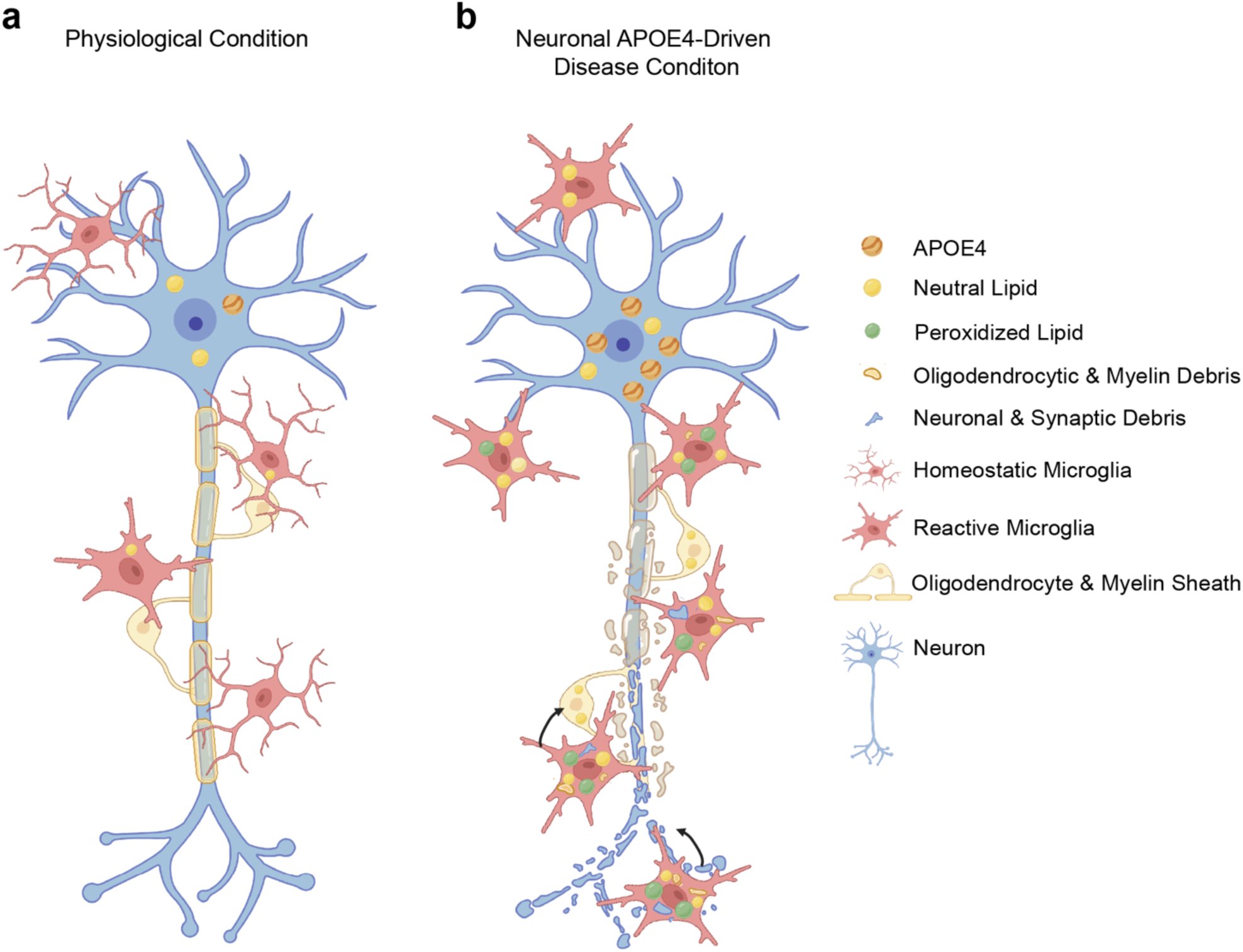
| Proposed model of neuronal APOE4-driven damaging lipid accumulation via contact-dependent neuron-oligodendrocyte-microglia interaction in Alzheimer’s disease (AD). **a,** Under physiological conditions, neurons, oligodendrocytes, and microglia maintain balanced lipid exchange to support neuronal activity and membrane turnover. Homeostatic levels of APOE promote efficient lipid trafficking and recycling, allowing microglia to clear and repurpose excess neuronal or myelin-derived lipids. Microglia remain in a homeostatic state, characterized by dynamic process motility, efficient phagocytosis, and minimal lipid droplet (LD) levels. **b,** In the presence of increased neuronal APOE4 under a stress condition, lipid homeostasis becomes disrupted. Stressed APOE4-expressing neurons exhibit increased production and release of lipids, overwhelming systematic lipid regulatory mechanisms. Microglia in contact with these neurons internalize excessive lipids through contact-dependent uptake, leading to accumulation of LDs, neutral lipids, and ROS induced damage. Concurrently, neuronal APOE4-induced stress and damage lead to oligodendrocyte lipid accumulation and dysfunction, contributing additional lipid burden to microglia via a contact-dependent process. The resulting feed-forward cycle of lipid accumulation, oxidative stress, and neuroinflammation propagates neuronal injury and circuit vulnerability. Together, this model positions neuronal APOE4 as an upstream driver of lipid dyshomeostasis that orchestrates neuron–oligodendrocyte-microglia metabolic coupling and contributes to AD progression.

